# Antifungal tolerance and resistance emerge at distinct drug concentrations and rely upon different aneuploid chromosomes

**DOI:** 10.1101/2022.11.30.518455

**Authors:** Feng Yang, Eduardo FC Scopel, Hao Li, Liu-liu Sun, Nora Kawar, Yong-bing Cao, Yuan-Ying Jiang, Judith Berman

## Abstract

Antifungal drug tolerance is a response distinct from resistance, in which cells grow slowly above the minimum inhibitory drug concentration (MIC). Here we found that the majority (69.2%) of 133 *Candida albicans* clinical isolates, including standard lab strain SC5314, exhibited *temperature-enhanced* tolerance at 37°C and 39°C, and were not tolerant at 30°C. Other isolates were either *always* tolerant (23.3%) or *never* tolerant (7.5%) at these three temperatures, suggesting that tolerance requires different physiological processes in different isolates. At supra-MIC fluconazole concentrations (8-128 μg/ml), tolerant colonies emerged rapidly at a frequency of ~10^−3^. In liquid passages over a broader range of fluconazole concentrations (0.25-128 μg/ml), tolerance emerged rapidly (within one passage) at supra-MIC concentrations. By contrast, resistance appeared at sub-MIC concentrations after 5 or more passages. Of 155 adaptors that evolved higher tolerance, all carried one of several recurrent aneuploid chromosomes, often including chromosome R, alone or in combination with other chromosomes. Furthermore, loss of these recurrent aneuploidies was associated with a loss of acquired tolerance, indicating that specific aneuploidies confer fluconazole tolerance. Thus, genetic background and physiology, and the degree of drug stress (above or below the MIC) influence the evolutionary trajectories and dynamics with which antifungal drug resistance or tolerance emerges.

**Importance:** Antifungal drug tolerance differs from drug resistance: tolerant cells grow slowly in drug, while resistant cells usually grow well, due to mutations in a few known genes. More than half of *Candida albicans* clinical isolates have higher tolerance at body temperature than they do at the lower temperatures used for most lab experiments. This implies that different isolates achieve drug tolerance via several cellular processes. When we evolved different strains at a range of high drug concentrations above inhibitory levels, tolerance emerged rapidly and at high frequency (one in 1000 cells) while resistance only appeared later at very low drug concentrations. An extra copy of all or part of chromosome R was associated with tolerance, while point mutations or different aneuploidies were seen with resistance. Thus, genetic background and physiology, temperature, and drug concentration all influence how drug tolerance or resistance evolves.

## Introduction

More than 1.5 million people die from invasive fungal infections every year (1). Increases in the global prevalence of fungal infections has become a major public health concern (2). This is largely because the at-risk population is expanding with the increase in patients with compromised immunity, who are especially vulnerable to fungal infection and the overall increase in lifespan in general. For the vast majority of fungal infections, high morbidity and mortality are caused by species from the genera *Candida, Aspergillus* and *Cryptococcus* species (1).

Only three antifungal drug classes are used clinically for monotherapy: the polyenes, azoles and echinocandins (3). Polyenes, such as amphotericin B, have potent and broad-spectrum antifungal activity and resistance is rarely seen. However, polyenes can cause severe toxicity in the kidneys and the liver (4), because they also bind to human cholesterol. Echinocandins are fungicidal against most *Candida* species and are first-line drugs for the treatment of candidemia (5). However, the relatively high cost of echinocandins and the need to administer them intravenously, makes them inappropriate in many settings. Azoles, such as fluconazole (FLC), inhibit 14α-lanosterol demethylase, a key enzyme in ergosterol biosynthesis. Azoles, which are fungistatic, have broad-spectrum antifungal activity, good safety profiles, relatively high bioavailability and are more affordable in many healthcare settings. Therefore, azoles are one of the most widely used antifungal drugs (6, 7).

The clinical prevalence of FLC resistance in *C. albicans*, a prevalent opportunistic human fungal pathogen, is generally less than 1% (8). Despite this, therapeutic failure of FLC against susceptible *C. albicans* isolates is often >30% in systemic infections (9). The fungistatic nature of azoles allows cells to survive and to evolve new traits, such as drug tolerance or resistance.

Since the introduction of first-generation azole drugs in the 1990s, most published research on drug responses did not distinguish between resistance and tolerance. Antifungal drug tolerance, which is distinct from resistance, has been best characterized in *C. albicans* cells responding to FLC ((10) and reviewed in (11)). Antifungal drug resistance is the ability to grow well at drug concentrations above a defined MIC for the drug. Antifungal drug tolerance is defined as a characteristic of drug-susceptible genotypes that can grow, albeit slowly, at inhibitory drug concentrations (12). Furthermore, in a tolerant isolate, not all cells in the isogenic population grow with similar dynamics. Furthermore, individual clinical isolates exhibit different levels of FLC tolerance, and the degree of tolerance changes as a function of growth conditions (9, 10). However, the degree to which tolerance in different isolates varies across environmental conditions remains to be characterized.

Mechanistically, antifungal resistance is usually due to genetic/genomic mutations that directly affect the drug-target interaction and these mutations affect the response of all cells in a population. Antifungal drug tolerance depends upon diverse stress pathway responses, including heat-shock responses, responses to amino acid starvation, kinases such as protein kinase C, and epigenetic processes (11). Stress pathways presumably enable the slow growth of some cells, and likely affect drug-target interactions indirectly. However, the mechanisms that affect tolerance and that distinguish tolerance and resistance are not well understood. Because the persistence and mortality of fungal infections is associated with high levels of tolerance (9, 10), we posit that antifungal tolerance, a property often overlooked in clinical assays, may explain at least some of the treatment failures caused by clinical isolates that test as susceptible. We also hypothesize that understanding how genetic and physiological processes that modulate tolerance should identify new strategies to improve the outcomes of antifungal drug therapies.

In this study, we first screened a collection of 133 clinical isolates to determine the prevalence of FLC tolerance under in vitro lab conditions at normal and at febrile body temperatures. We found three distinct types of temperature responses: 1) *temperature-elevated tolerance* (TET) at 37 and 39°C, relative to 30°C; 2) *always-tolerant* (AT) and never-tolerant (NT) isolates at the three temperatures. We then evolved representative TET and AT isolates for adaptation to FLC at a range of drug concentrations. We found that aneuploidy appears rapidly and is associated with increased tolerance, especially at drug concentrations above the MIC. By contrast, resistant isolates emerged at sub-MIC drug concentrations, only after ~5 days of passaging in sub-MIC drug concentrations and they proceeded to acquire higher resistance levels with time. Some of the resistant adaptors also acquired aneuploidies, but different ones from those in tolerant adaptors. This suggests that tolerance and resistance appear with distinct evolutionary dynamics and trajectories.

## Results

### Most clinical isolates tested were fluconazole-tolerant at physiologically relevant temperatures

Previously, we found that temperature and medium composition each affected antifungal tolerance to FLC and ketoconazole (10, 13). Therefore, we asked about how these two physiological factors affected one another. Interestingly, the effect of temperature on FLC tolerance was modulated, sometimes in opposite ways, by medium composition. For example, on RPMI-1640 and casitone plates, SC5314 was tolerant at *both* 30°C and 37°C, yet on chromagar, SD or SDC plates, the same strain was not tolerant at *either* 30°C or 37 °C. Interestingly, on YPD medium (a nutrient-rich medium), SC5314 had *temperature-enhanced* tolerance: it was *non-tolerant* at 30°C and *tolerant* at 37°C (Fig. S1). Thus, FLC tolerance is regulated by the interplay between temperature and medium composition.

To determine the role of temperature in tolerance across a broader set of clinical isolates, we surveyed 133 clinical *C. albicans* isolates for their susceptibility/resistance and tolerance levels in YPD medium under standard lab conditions (30°C) and at normal (37°C) and febrile (39°C) human body temperatures. We used disk diffusion assays analyzed by *diskImageR (14)* to measure the susceptibility/resistance (as the radius of the zone of inhibition (RAD_20_,)) and the tolerance (as the fraction of growth (FoG_20_,) in the zone of inhibition relative to growth outside it) for each strain. For most strains (69.2%), tolerance levels were affected by temperature, with higher tolerance levels at 37°C and 39°C than at 30°C, and little difference between tolerance levels measured at 37°C vs 39°C (Fig. 1A). The susceptibility/resistance levels remained similar at the three temperatures. Thus, tolerance, but not resistance, was affected by both medium composition and temperature.

**Figure 1.**
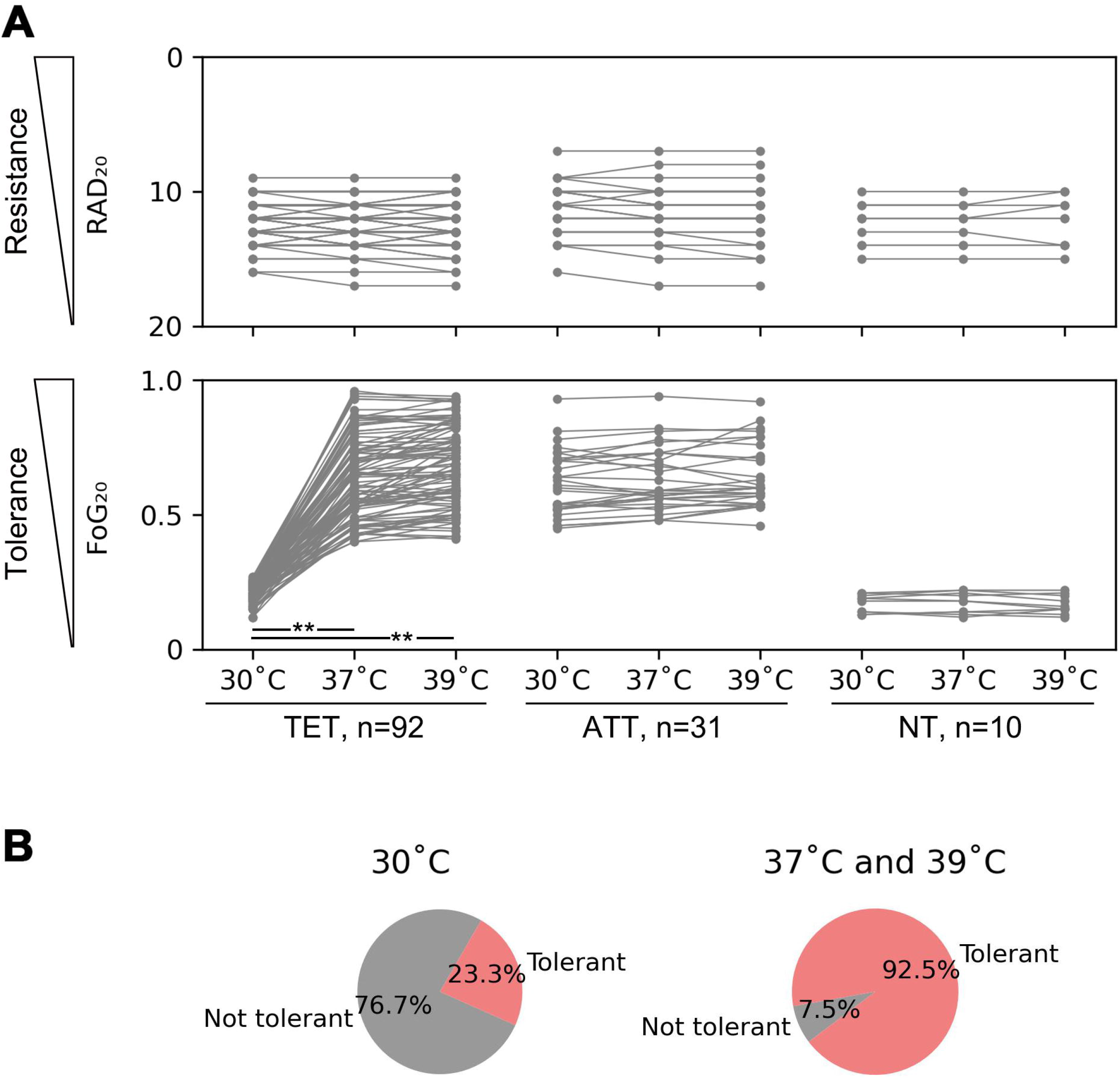
Measuring tolerance levels at 30°C, 37°C, and 39°C identifies three classes temperature response. Disk diffusion assays were performed on 133 different clinical isolates at the indicated temperatures (30°C, 37°C, and 39°C). The isolates were classified as temperature-enhanced tolerant (TET), all-temperature tolerant (ATT), and non-tolerant (NT), based on FoG_20_ values at the three temperatures. ** indicates p-value<0.001 as determined by two-tailed paired t-test (**A)**. **B** is the distribution of tolerant isolates at different temperatures (n=133). At 30 °C, 69.2% were TET, 23.3% ATT, 7.5% NT. Thus, the vast majority of strains exhibit tolerance at least at some temperatures.

We classified the isolates into three groups based on the effect of temperature on growth temperature-dependence of the tolerance (FoG_20_ values): <u>t</u>emperature-<u>e</u>nhanced <u>t</u>olerant (**TET),** (non-tolerant at 30 °C but highly tolerant at higher temperatures (n=92, average FoG_20_ values of 0.21 ± 0.03, 0.63 ± 0.17, 0.68 ± 0.15, at 30°C, 37°C and 39°C, respectively)); <u>a</u>ll-<u>t</u>emperature <u>t</u>olerant (**ATT**), which had similar FoG_20_ values at all three temperatures (n=31, FoG_20_ values of 0.52 ± 0.12, 0.61 ± 0.15, 0.61 ± 0.15, at 30°C, 37°C and 39°C, respectively); and <u>n</u>on-<u>t</u>olerant (**NT**), which exhibited only baseline levels of FoG_20_ at all three temperatures (n=10, average FoG_20_ values 0.21 ± 0.03, 0.23 ± 0.03, 0.24 ± 0.03, at 30°C, 37°C and 39°C, respectively). At 37°C and 39°C, 92.5% (123 out of 133) of the strains exhibited similar tolerance levels, while at 30°C, 76.7% (102 out of 133) were not tolerant on the YPD medium (Fig. 1B). Thus, both body and febrile temperature promoted tolerance on YPD medium in most isolates.

### Growth dynamics of tolerant cells are similar in a broad range of supra-MIC fluconazole concentrations

To better characterize the similarities and differences in the drug responses of the different classes of isolates, we used E-Test^®^ strips to measure susceptibility as the MIC after 24 h of growth, and spot dilution assays (13), analyzed at 48h, to measure tolerance. One representative isolate was used for each of the three temperature-response classes: SC5314 (the *C. albicans* lab strain) for TET, YJB-T1891, for ATT, and YJB-T490 for NT strain classes. The three isolates had the same MIC value (1 μg/ml) at 30°C, 37°C and 39°C on YPD (Fig. 2A)). However, spot assays revealed significant differences in the ability of individual colonies to grow in the presence of FLC (Fig. 2B). For the TET isolate at 30°C, and for the NT isolates at all three temperatures, no growth was evident above the MIC, indicating that they were not tolerant under these conditions. By contrast, the isolates that exhibited tolerance (the ATT isolate at all three temperatures, and the TET isolate at 37°C and 39°C) grew similarly at all the drug concentrations tested (up to 128 μg/ml). Thus, for TET and NT tolerant isolates, tolerant growth does not exhibit much concentration-dependence from 2-128 μg/ml FLC.

**Figure 2.**
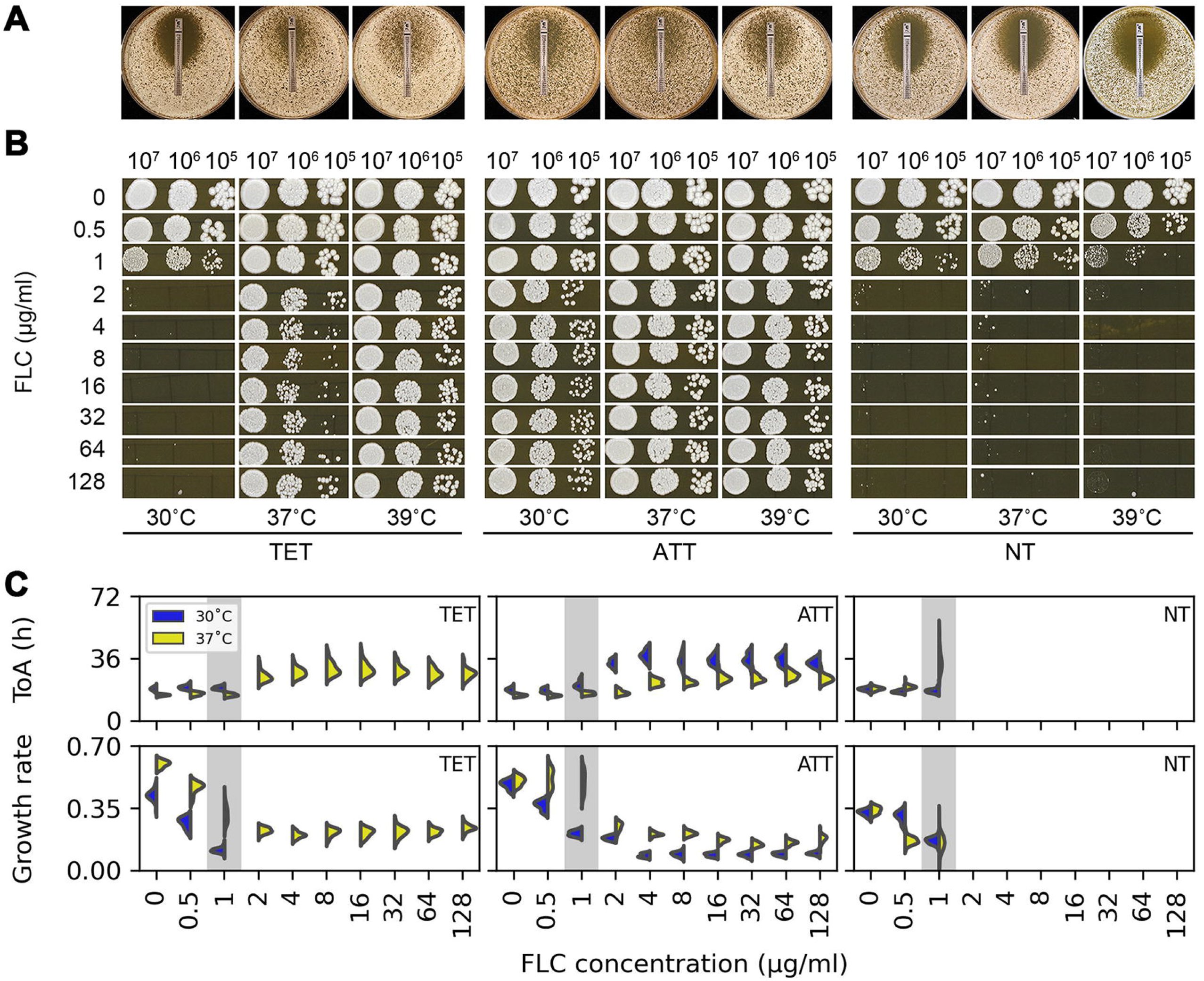
Temperature dependent growth dynamics of representative TET, ATT and NT strains in different drug concentrations. Representative isolates (TET, SC5314; ATT, YJB-T1891; and NT, YJB-T490) were assayed on plates with FLC E-Test strips (A) and on 10-fold dilution spot assays (B) at 30°C, 37°C and 39°C on YPD medium at the FLC concentrations indicated. In both cases, plates were photographed after 48h at the indicated temperatures. Growth dynamics of isolates in the presence of FLC were measured using *ScanLag* (10, 15). Approximately 100 colonies of the test strains were plated onto YPD plates supplemented with FLC at the concentrations indicated (C). The growth of colonies was monitored at 30°C and 37°C using *ScanLag*. Colony area was calculated from the area of light pixels in the images, a proxy for colony size, and the change in colony size over time was used to estimate colony growth rate. Note in (B) and (C), for tolerant isolates, growth was similar at all fluconazole concentrations equivalent to and above MIC. The graphs show the distribution of Time of colony Appearance (TOA, top panel) and growth rate (bottom panel) for all colonies on each plate, in 72 hours (top panel). The area of colonies occupied by light pixels and the change in this area over time are proxies for colony size and growth rate.

We investigated the impact of MIC and tolerance on population growth dynamics in the presence of FLC by plating 100-200 colony-forming units of each isolate on YPD plates supplemented with FLC at concentrations from 0.25-128 μg/ml. We monitored growth dynamics at both 30°C and 37°C for 48 h at 30 min intervals using *ScanLag*, which reports on the time of appearance and growth rates of individual colonies (10, 15).

At 30°C, colonies from TET isolate SC5314 failed to grow at FLC concentrations above the MIC (1 μg/ml), while at 37°C, growth was detectable. Colonies of the ATT isolate (YJB-T1891) grew at both 30°C and 37°C in all FLC concentrations, regardless of the MIC (1 μg/ml) (Fig. 2C) and for the NT isolate (YJB-T490), no colonies grew at either 30°C or 37°C in the supra-MIC FLC concentrations. These results are consistent with those from the E-test and spot dilution assays above.

*ScanLag* also measures colony growth on agar and reports the time required for a colony to become detectable on the plate. This time of appearance is considered a proxy of lag phase length (15). In all three strain types, at both 30°C and 37°C, the average time of appearance of colonies on drug plates was dose-independent at drug concentrations below the MIC. Above the MIC, the time of appearance of isolates that exhibited tolerance in the spot assays was longer than at sub-MIC drug concentrations, yet was dose-independent: the whole population appeared with a later time of appearance at all supra-MIC concentrations tested. In contrast to time of appearance, growth rate was dose-dependent at sub-MIC drug concentrations. Above the MIC, the growth rate, like the time of appearance, was slower than at sub-MIC concentrations, yet was drug dose independent. Thus, at supra-MIC drug concentrations, tolerant cells appeared later, presumably because they have a more prolonged lag phase, and grew slower than at sub-MIC concentrations. Nonetheless, these growth properties were similar at concentrations from 2 μg/ml to 128 μg/ml of fluconazole, consistent with the low degree of concentration-dependence of tolerance.

### Evolution of higher tolerance from cells in the non-tolerant state

To ask how a non-tolerant isolate adapts to supra-MIC FLC concentrations, we plated approximately 1×10^6^ cells of TET isolate SC5314 and NT isolate YJB-T490 on YPD plates supplemented with 8 μg/ml to 128 μg/ml FLC. The plates were incubated under conditions where these isolates were non-tolerant: 30°C for SC5314 and 30°C and 37°C for YJB-T490. As controls, we plated 1×10^6^ cells of SC5314 at 37°C and the ATT strain (YJB-T1891) at 30°C and 37°C on the same range of FLC concentrations. As expected, all controls produced a lawn of cells. By contrast, the TET strain at 30°C and the NT strain at both temperatures gave rise to a few hundred to a few thousand colonies (adaptors) per plate after 5 days on the drug plates (Fig. S2).

Disk diffusion assays were then performed on 90 randomly selected adaptors from SC5314 and YJB-T490 parent strains (18 adaptors from each 30°C drug plate). All these adaptors had a notable elevated tolerance (Fig. 3) when tested at 30°C. When tested at 37°C, all SC5314-derived adaptors (like the SC5314 ancestor) had high tolerance (Fig. 3 Top panel), while all YJB-T490-derived adaptors had low tolerance at 37°C (Fig. 3 Middle panel). Thus, adaptors derived from SC5314 at 30°C acquired tolerance at 30°C and maintained their tolerance at 37°C, and adaptors derived from YJB-T490 at 30°C acquired tolerance that was limited to 30°C. By contrast, many adaptors derived from YJB-T490 at 37°C were tolerant at both 30°C and 37°C (Fig. 3 Bottom panel). Thus, depending on the temperature used to evolve the adaptors, the NT isolate adapted to FLC in two distinct manners: at 30°C they acquired a new type of conditional tolerance that we term *<u>t</u>emperature-<u>s</u>ensitive* <u>t</u>olerance (**TST)**seen only at 30°C and not at 37°C; when evolved at 37°C, NT isolates acquired tolerance that was detectable at both 30°C and 37°C, a phenotype akin to that of other ATT-like adaptors.

**Figure 3.**
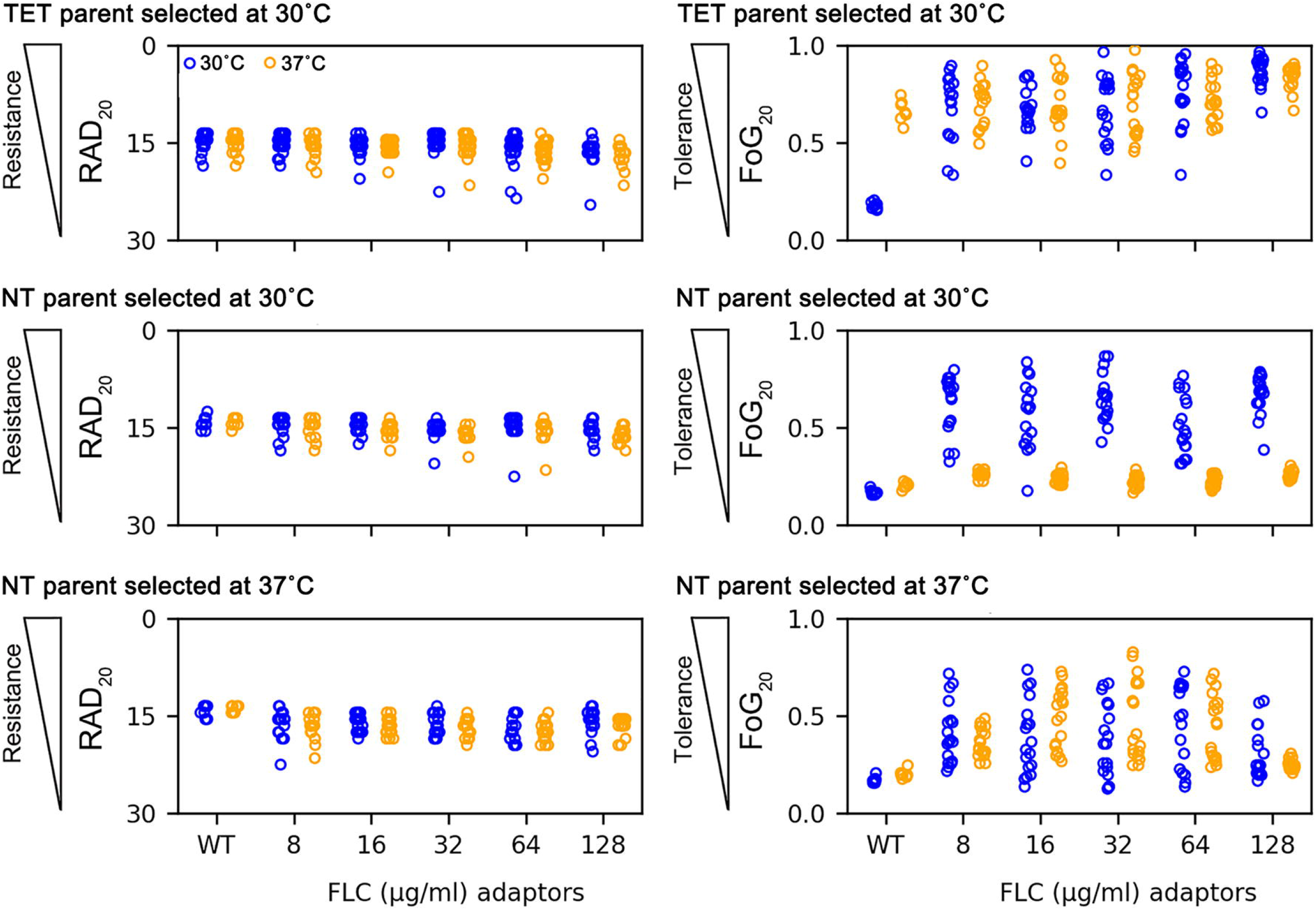
Selection for adaptors at supra-MIC fluconazole concentrations. FLC adaptors derived from TET strain SC5314 at 30°C, and those derived from the NT strain YJB-T490 at 30°C and 37°C, were tested for resistance (RAD_20_) and tolerance (Fog_20_) on disk diffusion assays at 30°C and 37°C. The disks contained 25 μg FLC. The plates were photographed at 24h and 48h for measuring RAD_20_ and FoG_20_, respectively. Each plot represents the data from 18 colonies that adapted to fluconazole concentrations indicated in YPD plates. In the parents, 16 individual colonies were tested.

### Recurrent aneuploidy enables the acquisition of fluconazole tolerance

To identify genomic mechanisms by which the SC5314 and YJB-T490-derived adaptors acquired tolerance, we performed deep sequencing of 18 independent adapted isolates for the TET and NT strains collected at 30°C on 8 μg/ml, 32 μg/ml, and 128 μg/ml of FLC. Taken together, 54 adaptors derived at 30°C from TET and NT isolates were sequenced.

Aneuploidy was prevalent among the adaptors. Out of the 54 TET-derived adaptors, one (FY9) that appeared on 8 μg/ml FLC plate grew poorly and was excluded from sequencing and further analysis. From the others, 50 (94.3%) were aneuploid (Fig. S3) and three were euploid. Among the 54 NT-derived adaptors at 30°C, 52 (96.3%) were aneuploid (Fig. S4) and two were euploid. Four of the five euploid adapters (from TET- and NT-derived) appeared at 8 μg/ml FLC, the lowest selective drug concentration tested. Variant calling of the 5 euploid adaptors revealed 11 high confidence *<u>de novo</u>* SNPs (1 to 4 per adaptor strain; 6 SNPs in the TET-derived adaptors and 8 SNPs in the NT-derived adaptors) (Table S1). Among them, a G2926C missense mutation of C1_06340W was detected in 4 of the 5 euploid adaptors; this encodes a protein of unknown function that has unphased allelic variation and contains ambiguous sequences (http://www.candidagenome.org/cgi-bin/locus.pl?locus=C1_06340W&organism=C_albicans_SC5314). Other mutations were all unique to a single adaptor: four of the remaining SNPs were in genes encoding proteins with unknown functions. None of the remaining 6 genes encodes any functions known to directly or indirectly affect drug responses (e.g., ergosterol biosynthesis, efflux pumps, lipid biosynthesis pathways or known stress response regulators). Mutation in one of the genes, *PGA63*, is similar to *S. cerevisiae SEC31*,which is annotated as having increased tolerance for aneuploid chromosomes (16), but the role of this gene in *C. albicans* will need to be investigated.

Analysis of adaptor karyotypes revealed several recurrent aneuploidies, several of which were seen in both the TET- and NT-derived isolates. The most prevalent adaptors (35 TET adaptors; 27 NT adaptors) had aneuploidy involving ChrR, either as trisomy of the whole chromosome (ChrRx3) (9 from TET, 10 from NT), monosomy of the left arm distal to the rDNA repeats (SegChrRx1) (6 from TET), or trisomy of ChrR from rDNA repeats to the right telomere (SegChrRx3) (12 from NT), or a combination of SegChrRx3 and SegChrRx1 (10 from TET). These ChrR aneuploidies appeared alone or in combination with trisomy of other chromosome(s) (Chr4 or Chr7 in TET adaptors; any combination of Chr4 and Chr6 trisomy alone or together with trisomy of Chr7 in NT adaptors) (Fig. 4). Five TET adaptors and 19 NT adaptors had aneuploidy of chromosome(s) other than ChrR. The 5 TET adaptors had a single additional aneuploidy: SegChr1×1, Chr4×3, Chr4×4, SegChr5×1, or Chr6×3, (Fig. S3). Among the 19 NT adaptors, only one had a single aneuploid chromosome (Chr5×1) and this monosomic strain grew very slowly. The remaining 18 adaptors had at least two aneuploid chromosomes, mostly Chr4×3 together with Chr6×3 (Fig. S3 and summarized in Fig. 4).

**Figure 4.**
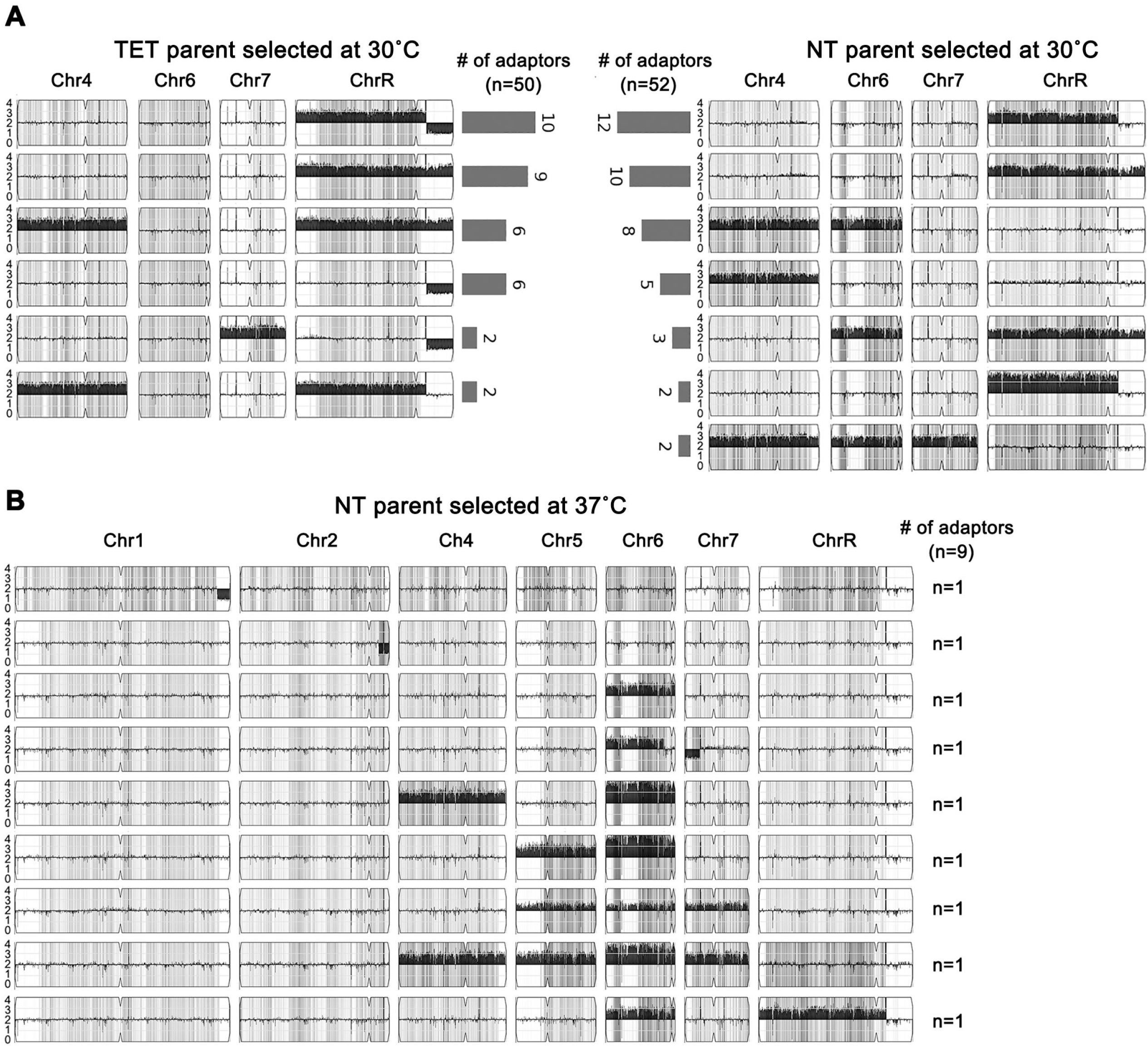
Recurrent aneuploidies associated with resistant and tolerant fluconazole adaptors. (A) Karyotypes recurrently identified in TET and NT derived fluconazole adaptors selected at 30°C. The total number of aneuploid adaptors, and the number of adaptors bearing each karyotype are also indicated. Chromosomes 1,2,3,5, which were euploid in these karyotypes, are not shown. (B) Karyotypes of 9 adaptors derived from NT isolate at 37°C. None of the karyotypes was identified in (A). The grey colored karyotypes include adaptors with aneuploidy of different homologs of the same chromosomes. The detailed karyotypes are shown in Fig. S3 and Fig. S4.

Taken together, NGS revealed 21 different karyotypes among the 50 TET-derived ATT-like aneuploid adaptors, and 17 different karyotypes among the 53 NT-derived TST aneuploid adaptors. Despite differences in genetic background and FLC concentration used for selection, the same aneuploidies (predominantly ChrRx3 or SegChrRx3) were recurrently associated with the emergence of FLC tolerance. This occurred in both the ATT types derived from TET ancestral isolates and the TST types derived from NT ancestral isolates.

The NT-derived adaptors obtained at 37°C also had diverse karyotypes. Each of the nine sequenced NT→ATT adaptors had a unique karyotype that was *not* seen in the NT→TST adaptors obtained at 30°C (Fig. 4): 7 had amplification of Chr6 alone (n=1) or in combination with aneuploidy (mostly trisomy) of one or more other chromosomes (n=6). Only two NT→ATT adaptors did not have Chr6 aneuploidy: one adaptor had SegChr1×1 alone, and one had SegChr2×1 alone.

Taken together, the karyotypes of the adaptors were FLC dose-independent but temperature-dependent. At 30°C, the TET- and NT-derived karyotypes involved recurrent ChrR amplification in TET→ATT and NT→TST adaptors selected across the range of FLC concentrations used. By contrast, from selection at 37°C, NT→ATT adaptors had distinctly different aneuploidy: predominantly Chr6 trisomy and tetrasomy, alone or in combination with another aneuploid chromosome. Thus, selection at 30°C yielded similar karyotypes with different tolerance temperature responses in different strain backgrounds. Moreover, different selection temperature (30°C vs 37°C) yielded different karyotypes in the same strain background (NT→TST vs NT→ATT) (Fig. 3 and Fig. 4).

### Loss of aneuploidy is associated with loss of acquired fluconazole tolerance

Whole chromosome aneuploidies can be gained and lost through mitotic missegregation. Previously we found that reversible chromosome gain and loss affected general fitness and drug responses in *C. albicans* (17, 18). Here, we asked if the FLC tolerance associated with recurrent aneuploidy was also unstable. Adaptors with different aneuploidies were spread onto YPD (without drug) and grown at 30°C for 36h. All yielded a mixture of colony sizes, with primarily small and some large colonies (Fig. 5A), although the frequency of appearance of large colonies was different for different aneuploids. We randomly selected one small and one large colony from each strain and tested them by disk diffusion assays at 30°C.

**Figure 5.**
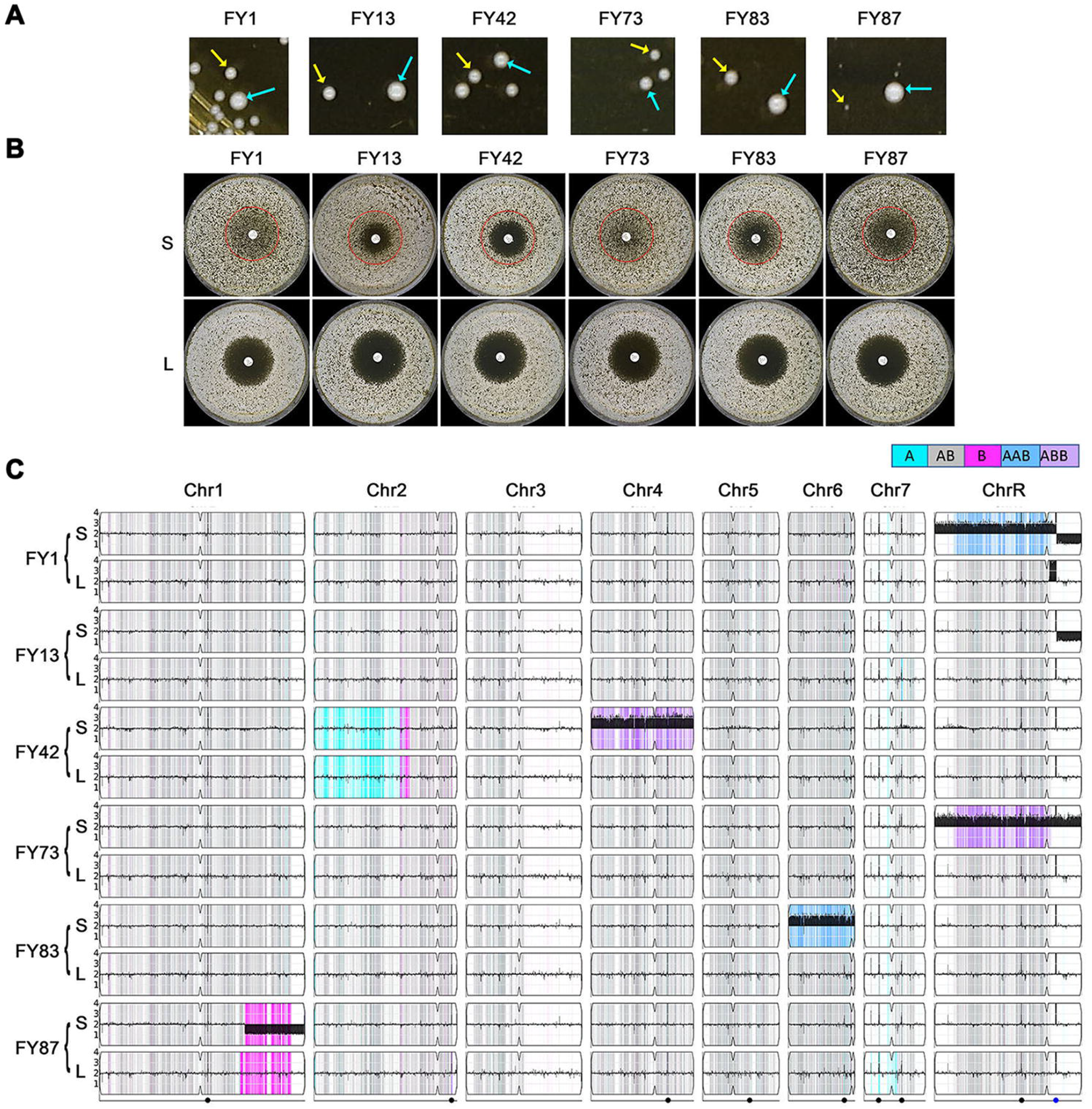
Tolerance acquisition and loss is associated with the gain and loss of aneuploid chromosomes. Strains with different aneuploid chromosomes were spread onto YPD plates and grown for 36 h at 30°C. Small colonies (S) and large colonies (L), indicated by yellow arrows and cyan arrows, respectively (A) were tested by standard disc diffusion assays (25 μg FLC per disk). The plates were incubated at 30°C for 48 h (B). Both S and L colonies were sequenced. The karyotypes were visualized by using Ymap (C).

To ask if point mutations had any role in the acquisition of tolerance, we performed variant calling to identify any SNPs of moderate to high effect (missense, nonsense, frameshift) that passed our quality filters (see Materials and Methods) in the small (aneuploid) and large colonies of the tolerant adaptors (Table S2). Small and large colonies from the same parent shared 14 SNPs. We assume that these SNPs were acquired during selection on FLC and were not drivers of tolerance. Taken together, in six tolerant adaptors representative of the six different karyotypes, we found variants that caused missense mutations in 23 genes. None of them were significantly (FDR > 0.1) enriched for gene ontology (GO) terms. Twelve genes were not annotated to any biological process, and 14 genes had unknown molecular functions. None of the SNPs found here were in genes encoding known or predicted drug efflux pumps or in known ergosterol biosynthesis enzymes or regulators.

Importantly, all the small colonies retained the higher tolerance seen in their parent strain <u>and</u> remained aneuploid like the adaptors from which they were derived. By contrast, all the large colonies were no longer tolerant (Fig. 5B) <u>and</u> had lost the aneuploid chromosome(s) to become euploid (Fig. 5C). This strong correlation, between the presence of aneuploidy and tolerance in all the small and loss of aneuploidy and tolerance in all the large progeny, supports the idea that the aneuploidies acquired during passaging on FLC were the major drivers of tolerance in the original adaptors. Because only some aneuploid chromosomes associated with tolerance, we presume that these aneuploidies appeared, in some cases recurrently, because they provide a selective advantage in the presence of drug (17, 19–23).

### Evolutionary trajectories differ at sub-MIC versus supra-MIC fluconazole concentrations

Because the TET and NT isolates acquired tolerance, but not resistance, when exposed to supra-MIC concentrations of FLC on solid medium, we next asked if this was a function of the FLC concentration. Therefore, we evolved one TET isolate (SC5314) and one NT isolate (YJB-T490) by passaging them daily (1:1000 dilutions) in YPD broth supplemented with FLC ranging from 4X below the MIC (0.25 μg/ml) to 128X above the MIC (128 μg/ml) for a total of 15 days. As a control, we also passaged the isolates in YPD without drug.

Every five days we randomly selected 18 or 96 colonies from the TET and NT cultures, respectively. In total, we measured 18 and 96 TET and NT derivatives per day (1, 5, 10, and 15 passages) and per drug concentration (0, 0.25, 0.5, 1, 2, 4, 8, 16, 32, 64 and 128 μg/ml FLC) for a total 792 (SC5314 derivatives) and 4224 (YJB-T490 derivatives) measurements, respectively (Fig 6A and 6B).

**Figure 6.**
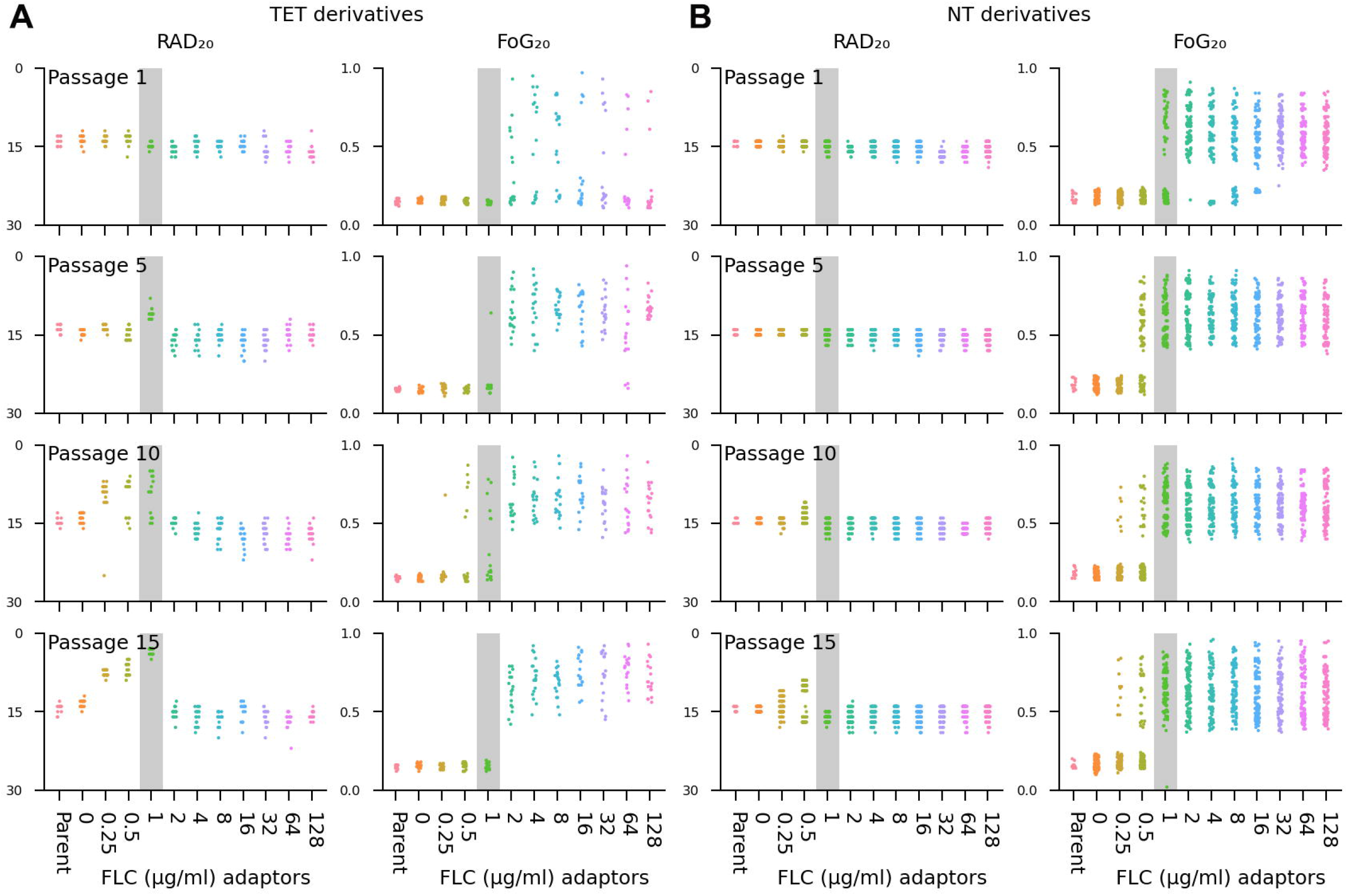
Distinct evolutionary trajectories associated with supra- and sub-MIC fluconazole concentrations. Emergence of tolerance (increased FoG_20_) and resistance (decreased RAD_20_) measured on disk diffusion assays for 18 isolates from each indicated passage of TET isolate SC5314 (A) and NT isolate YJB-T490 (B) for 1-15 days of propagation at the indicated drug concentrations. The MIC (1μg/ml) is highlighted in grey. Note that a broad range of tolerance emerges at all supra-MIC concentrations starting at passage 1, while increasing levels of resistance appear with time at or below the MIC only.

For SC5314 (MIC ~ 1 μg/ml FLC), among the 504 adaptors evolved at supra-MIC concentrations (2-128 μg/ml FLC), most exhibited tolerance (increased FoG_20_), while *none* of the adaptors acquired increased FLC resistance (Fig. 6A). In fact, some adaptors were *more susceptible* to FLC, as evidenced by a *larger* RAD_20_ than the parent strain. Similarly, for YJB-T490 (MIC ~ 0.75-1 μg/ml), most of the 2688 adaptors evolved at supra-MIC concentrations (2-128 μg/ml), exhibited tolerance, and *none* exhibited resistance (Fig. 6B). Interestingly, both the frequency with which tolerance arose and the range of tolerance levels achieved were similar at all the supra-MIC concentrations used for passaging (Fig. 6A). This suggests that once the drug inactivated its target, additional drug had little if any effect on how often or how much tolerance emerged.

At the MIC, SC5314 adaptors gained either resistance or tolerance (but not both) after 5 and 10 passages, while only resistant isolates appeared after 15 days of passaging at the MIC. Thus, it appears that at the MIC, SC5314 adaptors that were tolerant after 5 and 10 passages became resistant after 15 passages. Furthermore, there was stepwise increase in the level of resistance in adaptors evolved for 5, 10 and 15 days in 1 μg/ml FLC (Fig. 6A and Fig. S5). For YJB-T490 adaptors, only tolerant isolates were detected at the MIC. In some assays, the MIC of YJB-T490 is slightly lower than that of SC5314, and thus 1 μg/ml may be slightly higher than the actual MIC for this strain.

At sub-MIC concentrations (0.25 and 0.5 μg/ml), SC5314 adaptors became either resistant (decreased RAD_20_) or tolerant (increased FoG_20_) (but not both). Among the adaptors evolved in 0.25 μg/ml and 0.5 μg/ml FLC, resistance emerged in 35 and 30 adaptors, respectively, while tolerance was detected in 1 and 6 adaptors, respectively. The remaining adaptors appear to have had more transient tolerance that had been lost during growth in the absence of FLC. Adaptors evolved for 10 and 15 days in 0.5 μg/ml FLC also exhibited stepwise increases in resistance. YJB-T490 adaptors also acquired either resistance or tolerance (but not both). Among adaptors evolved in 0.25 μg/ml FLC, tolerance appeared after ten passages; yet after 15 passages in 0.25 μg/ml FLC for 15 days, ten were tolerant, and 23 were resistant. At 0.5 μg/ml, the proportion of resistant YJB-T490 adaptors relative to tolerant adaptors increased over time (0 vs 58; 29 vs 18, and 72 vs 24, for adaptors from passages 5, 10 and 15, respectively (Fig. 6). While this is consistent with the idea that isolates may have acquired tolerance in early passages and then become resistant in the later passages, we cannot be sure that the resistant adaptors are progeny of the prior tolerant ones.

The dynamics of the emergence of tolerance and resistance also differed considerably. In both parental strain backgrounds tolerance emerged more rapidly, within a single passage, while the first resistant adaptors appeared after 5 passages and only in SC5314 at 1 μg/ml FLC, while adaptors passaged in 0.25 μg/ml and 0.5 μg/ml only produced resistant and tolerant progeny after 10 days, and resistant adaptors after 15 days. In YJB-T490, *tolerant* adaptors evolved at or below the MIC 1 μg/ml, 0.5 μg/ml and 0.25 μg/ml initially appeared in passages 1, 5, and 10, respectively, while *resistant* adaptors that emerged from 0.5 μg/ml and 0.25 μg/ml FLC initially appeared on days 10 and 15, respectively. Thus, the acquisition of resistance and tolerance occurred at different relative stress levels (sub-MIC vs supra-MIC) and with different temporal dynamics (Fig. 6), and tolerant isolates generally appeared more rapidly than resistant isolates.

### Distinct and diverse genetic alterations in fluconazole-resistant versus fluconazole-tolerant adaptors

Individual colonies that arose following the evolution of SC5314 in broth passages had a range of resistance or tolerance levels, suggesting that they may have acquired distinct genetic or genomic changes. To test this hypothesis, we sequenced 73 randomly selected adaptors (34 resistant and 39 tolerant) derived from SC5314. Large-scale genome changes were visualized with Ymap (Fig. 7 and Fig. S6), and variant calling was performed to identify mutations in predicted open reading frames, focusing on missense mutations and frameshift mutations.

**Figure 7.**
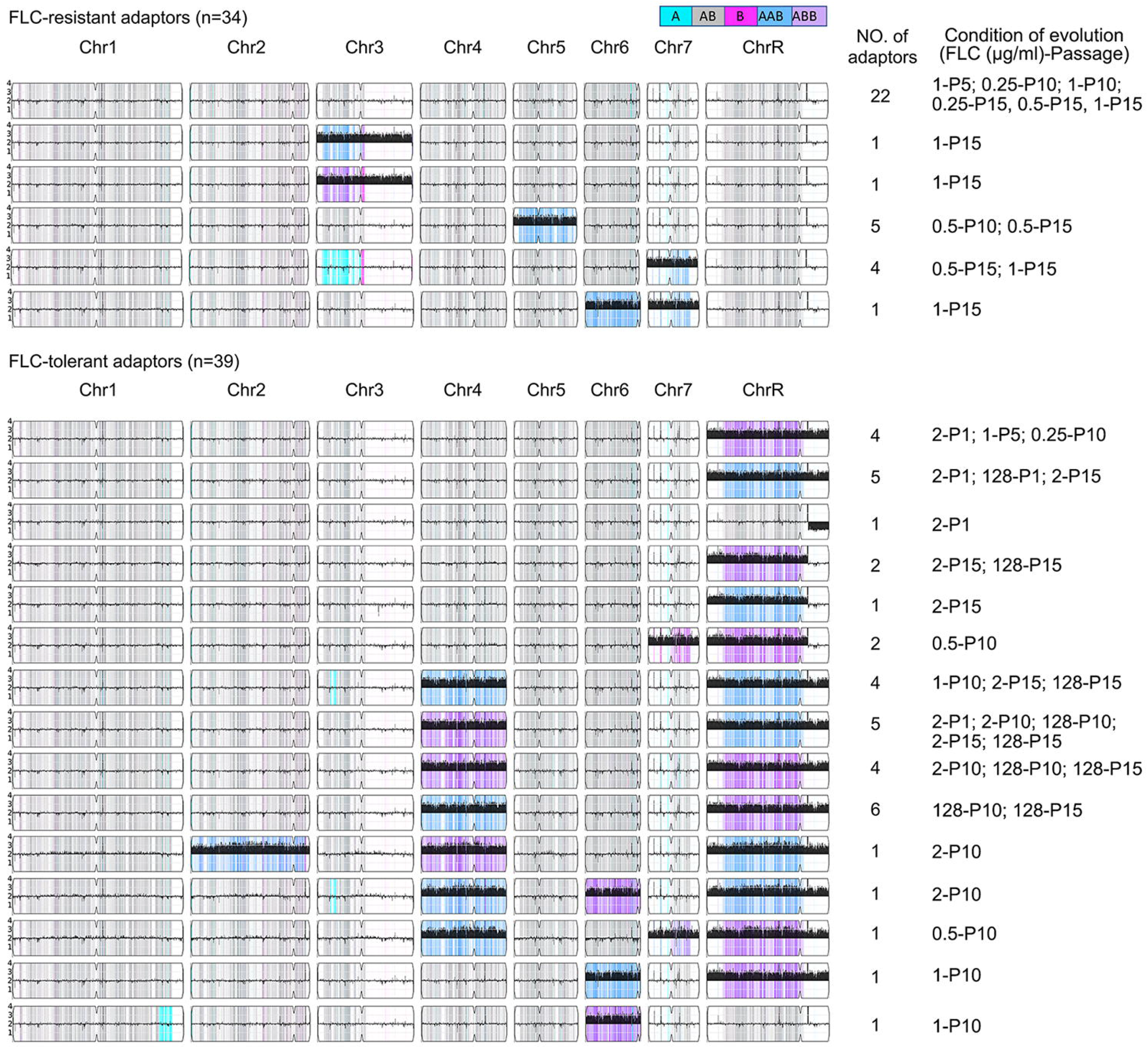
Distinct genomic changes in resistant and tolerant adaptors. DNA sequences of 34 FLC-resistant and 39 FLC-tolerant adaptors evolved from TET strain SC5314 were analyzed by Ymap (49). The karyotypes, number of adaptors with this karyotype. and the drug conditions (number of days evolved and FLC concentration) are indicated to the right of each karyotype diagram. For example, D5-1 indicates that the adaptors were evolved for 5 days in 1 μg/ml FLC. Colors indicate allele frequencies in the data as indicated in the key. Black histograms indicate the log1 ratio of DNA copy number for the strain indicated relative to a diploid control

Notably, all 39 tolerant adaptors that were sequenced were aneuploid (Fig. 7, lower section): 13 had whole or segmental aneuploidy only for all or segments of ChrR: (ChrRx3 (n=9), SegChrRx3 from 1.89Mb to right telomere (n=3), or SegChrRx1 from 1.89Mb to right telomere (n=1)). An additional 25 adaptors were aneuploid for ChrR in combination with aneuploidy of another one or two chromosomes (n=22 and n=3, respectively). Only one tolerant adaptor (Chr6×3, ABB, evolved over 10 passages in 1 μg/ml FLC) did not include copy number changes on ChrR. Thus, ChrR aneuploidy was the most prevalent aneuploidy associated with tolerance.

Among the 34 resistant adaptors sequenced, the majority (22/34) were euploid and 12 were aneuploid. Among the aneuploid adaptors, two (SY60 and SY61) had Chr3×3, five (SY20, SY21, SY22, SY51, SY55) had Chr5×3, four (SY50, SY56, SY58, SY59) had Chr7×3 (AAB), and one adaptor (SY57) had Chr6×3 (AAB) together with Chr7×3 (AAB). Notably, none of the resistant adaptors affected ChrR copy number and the karyotypes of the resistant adaptors were distinct from the tolerant adaptors (Fig. 7 and Fig.S5). Therefore, tolerant and resistant adaptors acquired distinct genomic changes: all tolerant adaptors were aneuploid, with ChrR aneuploidy as the predominant change while resistant adaptors were mostly euploid or were trisomic for either Chr3, Chr5, or Chr7.

Importantly, similar types of karyotypes were detected in SC5314 and YJB-T490 derived tolerant adaptors selected on supra-MIC FLC concentrations on agar plates (Fig. S3 and Fig. S4). Furthermore, the karyotypes of tolerant adaptors appeared in a dose-independent and time-independent manner (Table 1 and Fig. 7). For example, ChrRx3 aneuploidies appeared after one day in 2 μg/ml or 128 μg/ml FLC, after 5 days in 1 μg/m FLC, after 10 days in 0.25 μg/ml FLC, and after 15 days in 2 μg/ml FLC. Similarly, ChrRx3+Chr4×3 appeared after 1 day in 2 μg/ml FLC, 10 days in 1 μg/ml, 2 μg/ml and 128 μg/ml FLC, and after 15 days in 2 μg/ml and 128 μg/ml FLC. And segChrRx3 (from the left telomere to 1.89 MB) appeared after 10 days in 0.5 μg/ml FLC, and 15 days in 2 μg/ml and 128 μg/ml FLC (Table 1). Thus, the same aneuploidies can arise at different concentrations and at different times after initial drug exposure.

**Table 1.**
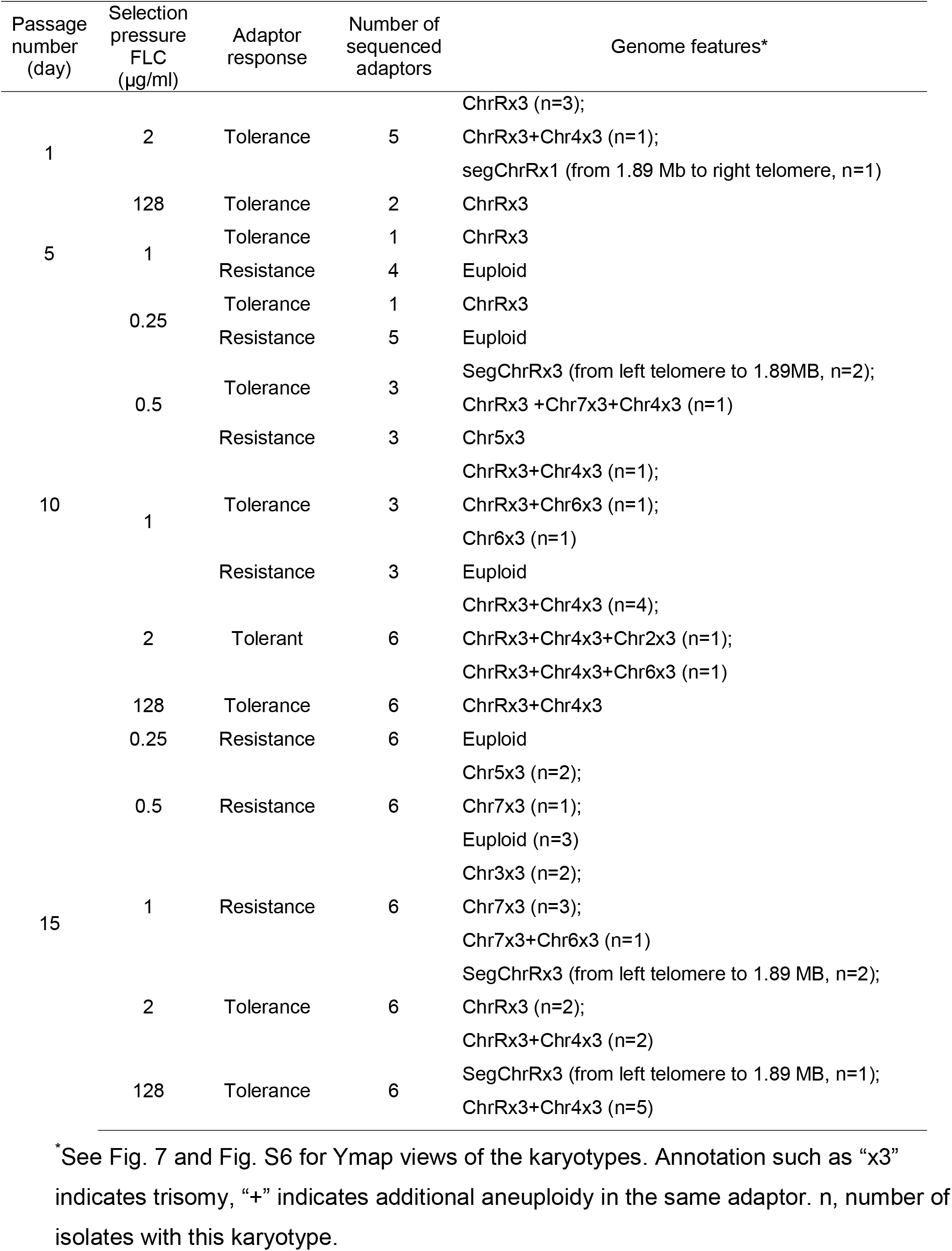
Karyotypes of the 34 resistant and 39 tolerant evolved progeny analyzed.

To ask about point mutations that may influence the degree of tolerance or resistance, we identified SNPs in the 39 tolerant and 34 resistant adaptors (Table S3). In total we found from 2 to 5 SNPs per tolerant isolate and 0 to 2 SNPs per resistant isolate. One of the two ORFs with detectable SNPs in both a tolerant and a resistant adaptor was C2_08380C, a possible ortholog of *S. cerevisiae DPB11*, that, when mutated, causes gross chromosomal rearrangements, chromosome loss and an increase in spontaneous DNA damage (24). While the degree to which these specific SNPs affect the encoded function is not known, this gene might contribute to increased frequencies of mutations in the adaptors.

SNPs found only in *tolerant* adaptors were detected in 27 ORFs. However, of these 27 only one could be associated with drug tolerance: C1_04010C. We detected 2 SNPs in this ORF in 4 different adaptors. This gene encodes a protein with a predicted NADP-dependent oxidoreductase domain, and its transcript is induced by ketoconazole, yet is repressed by Upc2p, a transcription factor that up-regulates ergosterol biosynthesis (25). There was no statistically significant (FDR > 0.1) gene ontology term enrichment of these 27 genes. Thus, the few SNPs that appeared may be neutral mutations and do not appear to have any obvious connections to drug resistance or tolerance.

SNPs found only in *resistant* adaptors were detected in 7 genes with no detectable gene ontology term enrichment (FDR > 0.1), but none were in genes that encode efflux pumps, or that are known to affect ergosterol biosynthesis, the two major mechanisms of azole resistance. We suggest that some of these SNPs are probably neutral or nearly-neutral mutations that arose during the course of passaging and/or false positive variants that our quality filters were not stringent enough to filter out.

## Discussion

Contrary to the concept from bacteria that antimicrobial drugs primarily select for the rapid evolution of drug resistance, here we found that *C. albicans*, a common cause of systemic bloodstream infections, primarily acquired tolerance to the widely used drug FLC. Importantly, this antifungal tolerance appeared rapidly, in some cases within a single day of exposure to drug concentrations above the strain MIC, while resistant isolates appeared after 5 days of exposure to low levels of FLC. Furthermore, the resistant isolates exhibited a stepwise increase in MIC levels (Fig. 6A), and we found no evidence that mutations in classic genes associated with FLC resistance played any role in the resistance acquired under the conditions used.

Here, we found that most clinical *C. albicans* isolates exhibit FLC tolerance at physiologically relevant temperatures, and not at 30°C, the temperature often used in lab studies (Fig. 1). This antifungal tolerance is largely dose-independent at supra-MIC drug concentrations (Fig. 2). Experimental evolution in supra-MIC drug concentrations rapidly selected for the acquisition of tolerance that was associated with a recurrent set of aneuploidies (Figs. 3,4,6,7). Resistant isolates only appeared at or below the MIC, after 5 or more days of passaging (Fig. 6). While most resistant isolates were not aneuploid, those that carried recurrent aneuploidies had extra copies of different chromosomes from those seen in tolerant isolates (Fig. 7). Furthermore, resistance (but not tolerance) appeared to be acquired in a stepwise manner (Fig. 6). Thus, experimental evolution at sub-MIC FLC concentrations selected for isolates that were either only tolerant or only resistant, and a different set of recurrent aneuploidies conferred tolerance vs resistance.

Of the original 133 clinical *C. albicans* isolates tested at 37°C and 39°C, up to 92.5% of the isolates were tolerant, while at 30°C only 23.3% of isolates were tolerant. Thus, it appears that FLC tolerance is prevalent in clinical isolates, at least those collected from Israeli patients. These clinical strains are largely euploid; thus, we suggest that tolerance is primarily due to genomic factors, including inherent genetic backgrounds, that modulate stress responses in different physiological conditions, such as temperature and medium composition. In addition, the acquisition of aneuploidy can increase or modulate tolerance levels. While this study focused on 133 Israeli clinical isolates, it will be interesting to determine if other clinical isolates exhibit temperature-enhanced tolerance.

Tolerant isolates were able to grow at supra-MIC FLC concentrations up to 128 μg/ml. Furthermore, the time of colony appearance, as well as growth rate of colonies on drug plates were dose-independent at supra-MIC FLC concentrations. Yet, below the MIC, colony growth rates were dose-dependent with colonies growing more slowly with increasing drug concentrations. This suggests that *C. albicans* mounts distinct cellular responses at sub-MIC versus supra-MIC drug concentrations. The concentration-independent nature of tolerant cell growth rates and the initial time of colony appearance at supra-MIC, and the concentration-dependent nature of resistance (Fig. 2B and C) at sub-MIC concentrations, highlights probably mechanistic differences between these processes. We posit that tolerance involves cellular stress mechanisms that are less sensitive to the intracellular drug concentration and enable growth when the drug target, in this case lanosterol demethylase, is completely inactivated by drug. The fact that different aneuploidies or mutations accompany tolerance vs resistance further supports this idea. Furthermore, isolates with different genetic backgrounds at the temperatures used and on agar or in broth media, adapted to supra-MIC FLC primarily by acquiring the same aneuploidies involving ChrR.

All but five tolerant adaptors had at least one aneuploid chromosome. In addition to the recurrent karyotypes involving ChrR aneuploidies, some tolerant adaptors acquired other aneuploidies. This suggests that individual cells use a similar strategy, chromosome instability, to reach non-identical outcomes (e.g., different levels of acquired tolerance) from different combinations of aneuploid chromosomes that provide a growth advantage in supra-MIC drug concentrations. Even though we detected a few SNPs in the aneuploid isolates and we do not know the degree to which they may (or may not) contribute to tolerance, the fact that loss of the aneuploid chromosome(s) was accompanied by loss of FLC tolerance suggests that aneuploidy was the primary mechanism conferring tolerance. The mechanism(s) by which the 5 euploid adaptors became FLC tolerant are less clear as the few SNPs identified in these strains did not encode obvious mediators of drug efflux, stress response or alternative strategies to bypass the deleterious effects of ergosterol depletion due to FLC exposure. While the SNPs or some other unidentified genetic change can not be ruled out, another possibility is that non-genetic metabolic shifts contributed to the ability of these isolates to grow despite the presence of supra-MIC drug concentrations (26).

Strain background and environmental conditions also contribute to the selective conditions that favor the acquisition of different aneuploidies. Interestingly, in both the TET and NT backgrounds, amplification of portions of ChrR enabled tolerance at 30°C but not at 37°C. Yet, for the NT strain, Chr6 trisomy increased tolerance at both 30°C and 37°C in the presence of the same range of FLC concentrations. Thus, the rapid emergence of tolerance can occur via multiple routes, and the trajectories of the routes are affected by the original strain background and the selection conditions. Future work will require analyzing the trajectories of adaptation of additional TET and NT isolates and identifying the chromosomal region(s) that specifically affect tolerance at different temperatures.

Some of the tolerant adaptors also became less FLC resistant (more susceptible, larger RAD_20_,). This was also seen for isolates passaged in posaconazole, another fungistatic azole (27). This supports the idea that the acquisition of antifungal resistance and antifungal tolerance follow different trajectories and that supra-MIC concentrations preferentially and can rapidly select for azole tolerance along with decreased azole resistance.

All tolerant strains acquired aneuploidy together and had a similar range of tolerance levels at all supra-MIC concentrations. By contrast, the resistant mutations that appeared at concentrations at or below the MIC, continued to become more resistant over time (Fig. 6 and Fig. S5). Interestingly, this stepwise acquisition of improved fitness was seen both with aneuploid and euploid resistant adaptors. This highlights the very different evolutionary trajectories that occur above and below the MIC. At all the supra-MIC concentrations tested, when the drug target is completely inhibited/saturated by drug, cells must respond with strategies independent of the drug-target interaction. By contrast, at sub-MIC concentrations, the drug target is not saturated, all cells are able to grow to some degree, providing the opportunity for the acquisition of a series of mutations that incrementally increased resistance. At sub-MIC, tolerance also emerged occasionally.

Why does aneuploidy appear so rapidly and with such a high frequency (~10^−3^) following exposure to supra-MIC drug concentrations? We suggest that three forces drive the rapid appearance of aneuploidy at high frequency: first, random aneuploid chromosomes maintained in the parent population may provide standing variation; second, FLC induces chromosome mis-segregation via cell cycle defects that yield tetraploid intermediates and, subsequently, random aneuploids (28, 29) at high frequency; and third, strong selection for specific aneuploidies that provide an adaptive advantage in the presence of the drug allows the aneuploid isolates to compete and outgrow arrested euploid cells (18). Chromosome mis-segregation events that lead to whole chromosome aneuploidy are relatively frequent (every 5×10^5^ cell divisions in yeast (30) and on the order of once every 10^4^ to 10^5^ divisions in mammalian cells (31)). Since *C. albicans* chromosomes contain between 407-1383 genes per chromosome (http://www.candidagenome.org/cache/C_albicans_SC5314_genomeSnapshot.html, as of Nov 24, 2022), it follows that a single aneuploid chromosome should affect the stoichiometry of many proteins. Since segmental aneuploidies, which are dependent upon recombination events, also appeared, it is tempting to speculate that either drug stress increases the likelihood of mitotic recombination as was seen for LOH events (32), or that recombination events are more frequent in strains carrying aneuploid chromosomes. Consistent with this idea, haploid *S. cerevisiae* lab strains carrying single extra chromosomes conferred increased mitotic recombination as well as chromosome instability (33).

In this study, tolerant adaptors emerged more rapidly (within one passage) than resistant adaptors. Ultimately, all supra-MIC isolates became tolerant, while only some of the sub-MIC adapters became resistant. Indeed, even resistant adaptors that had acquired resistance via aneuploidy appeared much later in the passaging and may have arisen from tolerant intermediates (e.g., some adaptors evolved in 0.5 μg/ml and 1 μg/ml FLC were tolerant on day 10, but all were resistant on day 15, and some were aneuploid (Fig. 6)). Thus, while the dynamics of acquiring tolerance is different at sub- and supra-MIC drug concentrations, a similar set of aneuploidies confers resistance, and a different set of aneuploidies can confer tolerance.

In *C. albicans*, two major mechanisms contribute to FLC resistance: alteration of the drug target and increased drug efflux. A combination of these mechanisms causes stepwise development of FLC resistance ((34); reviewed in (35)) or can arise in a single segmental aneuploidy (19). Surprisingly, we did not find any of the classic resistance mutations in resistant adaptors. This could be due to the high fitness cost of mutations in these genes (34, 36, 37). In the presence of sub-MIC FLC, cells are under weak stress and mutations with a high fitness cost might be outcompeted. In general, gain-of-function mutations in *MRR1, TAC*, or *UPC2* in FLC-resistant clinical isolates incur a fitness cost and are outcompeted by the matched susceptible isolates both *in vitro* and in *vivo* (36). Artificially introducing resistance-enhancing mutations causes a stepwise increase in resistance that is associated with a gradual reduction in fitness both in *vivo* and *in vitro* (34). Similarly, experimental evolution with increasing amounts of FLC yielded resistant mutants with mutations in *TAC1, UPC2*, and/or *ERG11* that were less fit than their parents in macrophages, as well as in the presence of several stresses, including cell wall stress, cell membrane stress, salt stress, oxidative stress and temperature stress (37). The longer time required for resistant adaptors to arise, even if aneuploid, provides indirect support for the idea that resistance is due to the accumulation of multiple mutational events.

Paradoxically, *C. albicans* strains that we surveyed from the clinic are euploid, not aneuploid, despite the observation that all adaptors that rapidly acquired increased tolerance were aneuploid. This implies that multiple changes in chromosome copy number can confer rapid increases in tolerance levels. Under drug selection, the aneuploidies should provide a fitness advantage and should be maintained at least until other, more stable mutations, with improved fitness, out-compete the aneuploids (38). Future experiments will address whether tolerant adaptors that acquired aneuploidies eventually evolve to lose the aneuploidy and retain high levels of drug tolerance and/or resistance.

Cells appear to adopt different adaptive trajectories at supra-MIC versus sub-MIC drug concentrations. At supra-MIC FLC, extreme stress may favor the immediate appearance of tolerance, perhaps because aneuploidy can appear within a single cell division or because aneuploidy may slow growth and thus might slow drug metabolism. At sub-MIC FLC concentrations, cells experience only mild stress that does not affect cell survival, which may enable the evolution of resistance in a stepwise manner. The degree of resistance, measured by the MIC or RAD, can increase over time of exposure to the drug (Fig. 6 and Fig. S5). Accordingly, more extended passaging at sub-MIC FLC concentrations yields adaptors with higher MIC levels. We speculate that this may involve a sequential acquisition of several different types of resistance mutations, including point mutations that confer stable drug resistance.

In bacteria, both sub- and supra-MIC antibiotic concentrations select for resistant mutants, but different selection strengths confer different evolutionary trajectories and drive the acquisition of different mutations (reviewed in (39). When selected in cidal antibiotics at supra-MIC drug concentrations, resistance mutations in one or a small number of “classical” resistance genes are selected. But when exposed to low antibiotic concentrations, bacteria accumulate mutations of several “non-classic” genes that individually confer weak resistance but together can confer higher resistance levels (40). Selection in subinhibitory concentrations also increase rates of mutation, recombination, and horizontal gene transfer, thereby increasing the appearance of genetic changes. In addition, selection at sub-MIC antibiotic concentrations can select for plasmids carrying multiple resistance genes that lead to multidrug resistance (MDR) (41).

In summary, FLC tolerance is prevalent in *C. albicans* clinical isolates, especially at 37°C and it exhibits different temperature responses in different strain backgrounds. The acquisition of resistance and tolerance occurs via distinct evolutionary trajectories that are largely a function FLC stress levels: supra-MIC FLC stress drives tolerance, which appears rapidly and enables growth at wide range of supra-MIC drug concentrations; sub-MIC FLC stress selects for either tolerance or stepwise elevated resistance. Different strains have different intrinsic levels of tolerance, but they generally appear to rapidly acquire additional tolerance by becoming trisomic for specific chromosomes. The most commonly acquired aneuploidy in tolerant adaptors is part or all of ChrR, while trisomy of Chr3 or Chr5, are associated with the relatively rapid appearance of increased antifungal drug resistance.

## Materials and Methods

### Strains and growth conditions

Strains used in this study are listed in Table S4. Stock cultures of all strains were preserved in 35% glycerol and maintained at −80°C. The 133 clinical *C. albicans* were collected from hospitals in Israel and kindly provided by Ronen Ben-Ami. Unless otherwise specified, cells were routinely grown on Yeast extract-Peptone-Dextrose (YPD)-agar media (1% [w/v] yeast extract, 2% [w/v] peptone and 2% [w/v] D-glucose, 2%[w/v] agar) at 30°C. Other media used in this study include casitone agar plates (0.9%[wt/vol] casitone, 0.5% [wt/vol] yeast extract, 1.15% sodium citrate dihydrate [wt/vol], 2% [wt/vol] glucose, 2% [wt/vol] d-glucose, and 2% [wt/vol] agar), SD agar plates (0.67% [wt/vol] yeast nitrogen base without amino acids, 2% [wt/vol] d-glucose, and 2% [wt/vol] agar), and SDC agar plates (0.67% [wt/vol] yeast nitrogen base without amino acids, 2% [wt/vol] d-glucose, 0.2% [wt/vol] complete amino acid mixture, and 2% [wt/vol] agar). Drugs were dissolved in dimethyl sulfoxide (DMSO) and stored at −20°C.

### E-test strip assays

E-tests were performed as described previously (20). Strains were streaked from −80°C freezer onto YPD agar. After incubation at designated temperature (30, 37 or 39°C) for 24h, colonies were chosen randomly and suspended in distilled water. Cell density was determined using a hemocytometer. Cells were adjusted to 1×10^6^ cells/ml, and 100 μl of the culture was spread on YPD agar. The plate with an FLC E-test strip (BioMerieux, Marcy l’Etoile, France) at the center was incubated at designated temperature for 24h and then photographed.

### Disk diffusion assays

Disk diffusion assays were performed as previously described (10, 13). The CLSI M44-A2 guidelines for antifungal disk diffusion susceptibility testing were followed with slight modifications. Briefly, strains were streaked from glycerol stocks onto YPD agar and incubated for 36h at designated temperatures. Colonies were suspended in distilled water and adjusted to 1×10^6^ cells/ml. 100 μl of cell suspension were spread onto 15□ml YPD plates. An empty paper disk (6□mm diameter, and 0.7 mm thickness) supplemented with 5 μl of 5 mg/ml FLC was placed in the center of each plate. Plates were then incubated at designated temperature and photographed at 24h and 48h. Analysis of the disk diffusion assay was done using the *diskImageR* pipeline (14). The fraction of growth inside the zone of inhibition and radius of inhibition, referred to as FoG and RAD throughout the manuscript, represent parameters measured at 20% drug inhibition (FoG_20_ and RAD_20_, respectively).

### Spot assay

Cells were suspended in distilled water and counted using a hemocytometer. Cell density was adjusted to 1×10^7^ cells/ml. 3 μl of 10-fold serial dilutions were spotted on YPD plates with or without drugs (control). The plates were incubated at the designated temperature and photographed after 2 days.

### ScanLag assay

The *ScanLag* assay was performed as described in (10) with minor modifications. Approximately 100 cells were spread onto YPD plates with or without FLC. The plates were placed on the scanners at designated temperature and scanned every 30 min for 48 h. Image analysis was done in MATLAB using the modified “ScanLag” script (10, 15).

### Obtaining drug adaptors from plates

Cells were adjusted to 1×10^7^ cells/ml as described above. 100 μl of the culture were spread on YPD plates supplemented with drugs. The plates were incubated at the designated temperature for 5 days. Adaptors were randomly chosen.

### Daily passage in YPD broth supplemented with fluconazole

SC5314 was inoculated into 1ml YPD broth at a final density of approximately 2.5×10^3^ cells/ml. The YPD broth was supplemented with DMSO (negative control) or with 2-fold increase of FLC from 0.25 μg/ml – 128 μg/ml FLC. Every 24h or when the OD600 of culture was higher than 1.0, 1 μl of each culture was inoculated into 1ml YPD broth supplemented with the same concentration of FLC. After 1, 5, 10, and 15 days, the cultures were washed and diluted with distilled water. Approximately 100 cells were spread on YPD plates. The plates were incubated at 30°C for 36 h and 18 colonies from each plate were randomly tested with disk diffusion assays using FLC-containing disks.

### Colony instability assay

As described previously (18), aneuploid strains were streaked from −80°C freezer to YPD plates and incubated at 30°C for 36 h. One small colony was randomly chosen and suspended in distilled water. Cells were diluted with distilled water and approximately 200 cells were spread on a YPD plate and incubated at 30°C for 36 h. One small (S) colony and one large (L) colony were randomly chosen for further studies.

### Next generation sequencing (NGS)

NGS was performed as described in (18).

### Variant calling

Paired-end short reads of all sequences obtained from NGS were trimmed with Trimmomatic (version 0.39; (42)) with default settings and read quality. Trimmed reads were aligned to both alleles of the *C. albicans* reference genome (GCF_000182965.3_ASM18296v3) with Burrows-Wheeler Aligner (bwa mem, version 0.7.17; (43)). The alignments were sorted with SAMtools (samtools view; samtools sort; version 1.15.1; (44)), and duplicates were marked and removed with Picard Tools (version 2.27.5; available at http://broadinstitute.github.io/picard/). Indels were then realigned with GATK (version 3.8-1-0; (45)), and the alignments were resorted and reindexed with SAMtools. Variants (SNPs and indels) were called using FreeBayes (-J - K --report-genotype-likelihood-max -a -F 0.1; version 0.9.21; (46)). Variants that were present in both ancestor and evolved strains were removed using BCFtools (isec; version 1.16-17; (44)), filtered for quality with BCFtools, and annotated with SNPeff (version 5.1d; (47)). The validity of each high-quality variant was then checked using the Integrative Genomics Viewer (IGV; version 2.15.1; (48))

### GO analysis

Gene Ontology Term Finder tool on Candida Genome Database (http://www.candidagenome.org/cgi-bin/GO/goTermFinder) was used for functional enrichment analysis, using default parameters. Only hits with p-value corrected for false positives FDR < 0.1 were considered as significant.

## Data availability

All sequence data are available in the ArrayExpress database at EMBL-EBI (www.ebi.ac.uk/arrayexpress) under accession numbers E-MTAB-12175, E-MTAB-12155, E-MTAB-12169 and E-MTAB-12189.

## Acknowledgements

We thank Jacob Steenwyk for helpful suggestions. This work was supported by the National Key Research and Development Program of China (2022YFC2303000), the National Natural Science Foundation of China (NO. 82020108032), and the Innovation Program of Shanghai Municipal Education Commission (NO. 202101070007E00094) to Yuan-ying Jiang. National Natural Science Foundation of China (NO. 81872910), and Shanghai Key Basic Research Project (NO. 19JC1414900) to Yong-bing Cao. Judith Berman also received funding from the European Research Council (ERC) under the European Union’s Horizon 2020 research and innovation programme (grant agreement No 951475).

## Supplementary materials

**Figure S1. Physiological factors affect fluconazole tolerance.**

*C. albicans* lab strain SC5314 was tested on disk diffusion assays with drug disks (25 μg FLC) on agar plates containing the indicated media and grown at the indicated temperatures. The plates were photographed at 48h to highlight tolerance (growth within the zone of inhibition.

**Figure S2. Growth of tolerant and non-tolerant strains on fluconazole plates.**

Approximately 1×10^6^ cells of TET (SC5314), ATT (YJB-T1891), and NT (YJB-T490) strains were grown on YPD plates supplemented with FLC at the indicated concentrations and incubated at 30°C and 37°C for 5 days. Estimated number of colonies on plates with 8 μg/ml~128 μg/ml drug plates are also shown.

**Figure S3. Karyotypes of all adaptors evolved from SC5314 at 30°C.**

SC5314-derived adaptors from YPD plates supplemented with 8 μg/ml, 32 μg/ml, and 128 μg/ml, were deep-sequenced and karyotype maps were generated by using the Ymap. Each chromosome is illustrated to scale. The centromere positions are indicated by an indentation. The major repeat sequence (MRS) positions are indicated by a black square below the chromosome. Local copy number estimates, scaled to the genome ploidy, are displayed as black histograms along the length of the chromosome. Allele frequencies are color coded: homolog “a” is cyan, homolog “b” is magenta and heterozygous alleles are grey. Karyotypes of all the adaptors sequenced are shown. The order of the adaptors is the same as the order when they were picked up from the fluconazole plates.

**Figure S4. Karyotypes of adaptors derived from NT strain YJB-T490 at 30°C.**YJB-T490-derived adaptors chosen from YPD plates supplemented with 8 μg/ml, 32 μg/ml, and 128 μg/ml at 30°C were deep sequenced. Karyotypes of all the 54 adaptors sequenced are shown. Homozygosis of alleles is indicated in red.

**Figure S5. Stepwise increase in MICs for resistant adaptors evolved in 1 μg/ml fluconazole.**

On day 5, day 10, and day 15, the MIC of one resistant adaptor from the 1μg/ml cultures was tested with FLC Etest strips. As a control, the parent and one adaptor isolated from passage 1 were also tested. The plates were incubated at 30°C for 48h and then photographed.

**Figure S6. Karyotypes of progeny evolved in YPD broth supplemented with fluconazole.**

34 FLC-resistant and 39 FLC-tolerant progeny were sequenced. FLC concentration and timepoint of test are indicated above each section. Strain name and whether it exhibited resistance (R) or tolerance (T) are indicated to the left of each karyotype.

**Table S1:**Mutations detected in coding sequences in 5 euploid fluconazole-tolerant adaptors.

**Table S2:**Mutations detected in coding sequences in 6 pairs of small and large colonies.

**Table S3:**ORFs having mutations identified in the 34 fluconazole-resistant and the 39 fluconazole-tolerant adaptors.

**Table S4:**Strains used in this study.

## Notes

### Competing Interest Statement

The authors have declared no competing interest.

### Summary of Updates

We added the variant analysis of the 5 euploid tolerant adaptors, and we edited the Discussion accordingly.

