## Supplemental Table S4 for "Antifungal tolerance and resistance emerge at distinct drug concentrations and rely upon different aneuploid chromosomes"

Table S4. Strains used in this study

| Strain | Karyotype | FLC tolerance | Selection |
| --- | --- | --- | --- |
| SC5314 | Euploid | TET^*^ | No selection |
| YJB-T490 | Euploid | NT | No selection |
| YJB-T1891 | Euploid | ATT | No selection |
| FY1-FY18 | See Fig. S3 | ATT | SC5314 derived FLC (8 μg/ml) adaptors |
| FY37-FY54 | See Fig. S3 | ATT | SC5314 derived FLC (32 μg/ml) adaptors |
| FY73-FY90 | See Fig. S3 | ATT | SC5314 derived FLC (128 μg/ml) adaptors |
| FY445-FY462 | See Fig. S4 | TST | YJB-T490 derived FLC (8 μg/ml) adaptors |
| FY481-FY498 | See Fig. S4 | TST | YJB-T490 derived FLC (32 μg/ml) adaptors |
| FY517-FY534 | See Fig. S4 | TST | YJB-T490 derived FLC (128 μg/ml) adaptors |
| SY1-SY73 | See Fig. S6 | Some are resistant. Some are tolerant | SC5314 daily passaged in 0.25~128 μg/ml |

^*^TET: temperature-enhanced tolerant NT: non-tolerant; ATT: all-temperature tolerant; TST: temperature-sensitive tolerant.
