## Supplemental figures S1-S6 for "Antifungal tolerance and resistance emerge at distinct drug concentrations and rely upon different aneuploid chromosomes"

### Figure S1

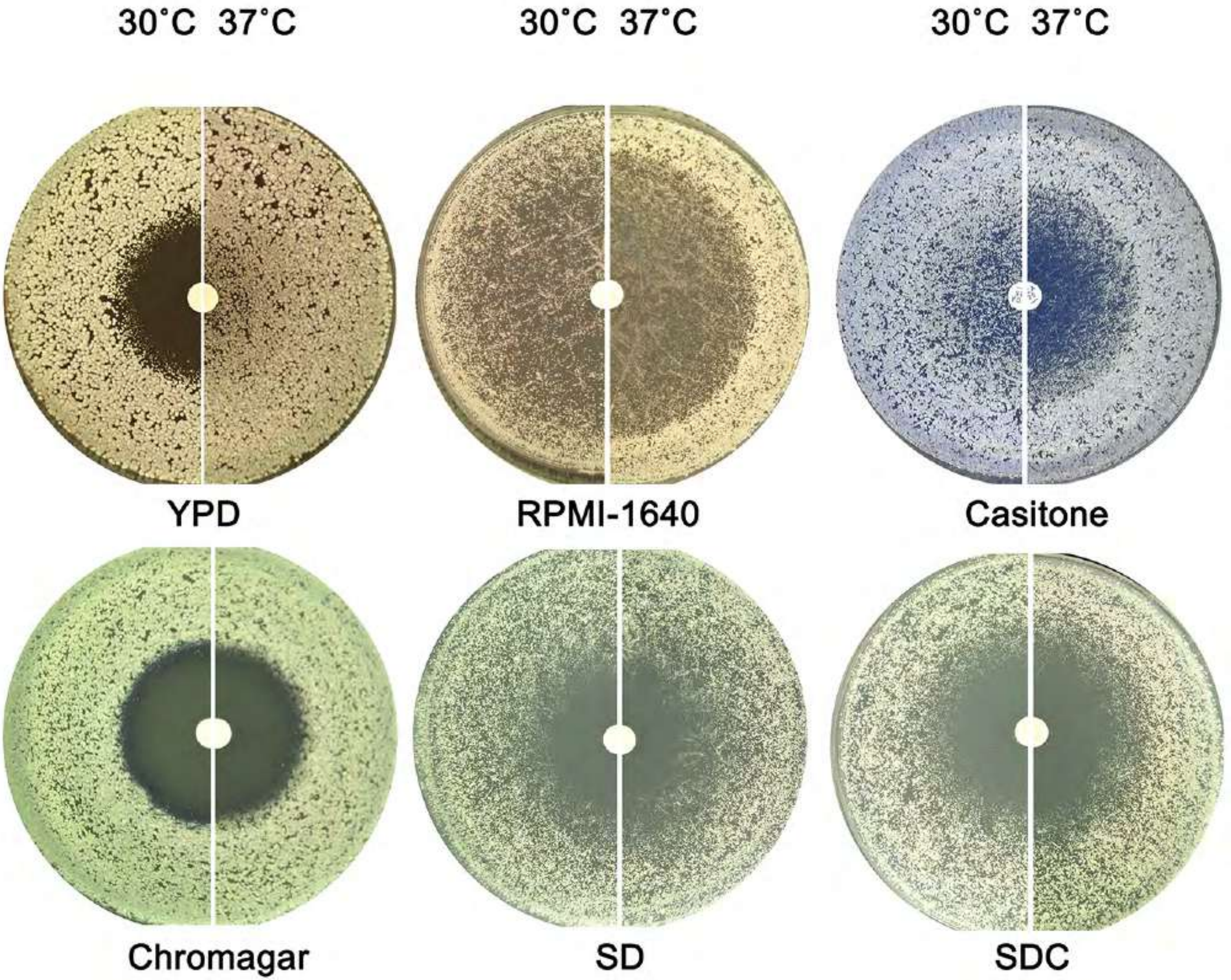
30˚C 37˚C
30˚C 37˚C
30˚C 37˚C
YPD
RPMI-1640
Casitone
Chromagar
SD
SDC

### Figure S2

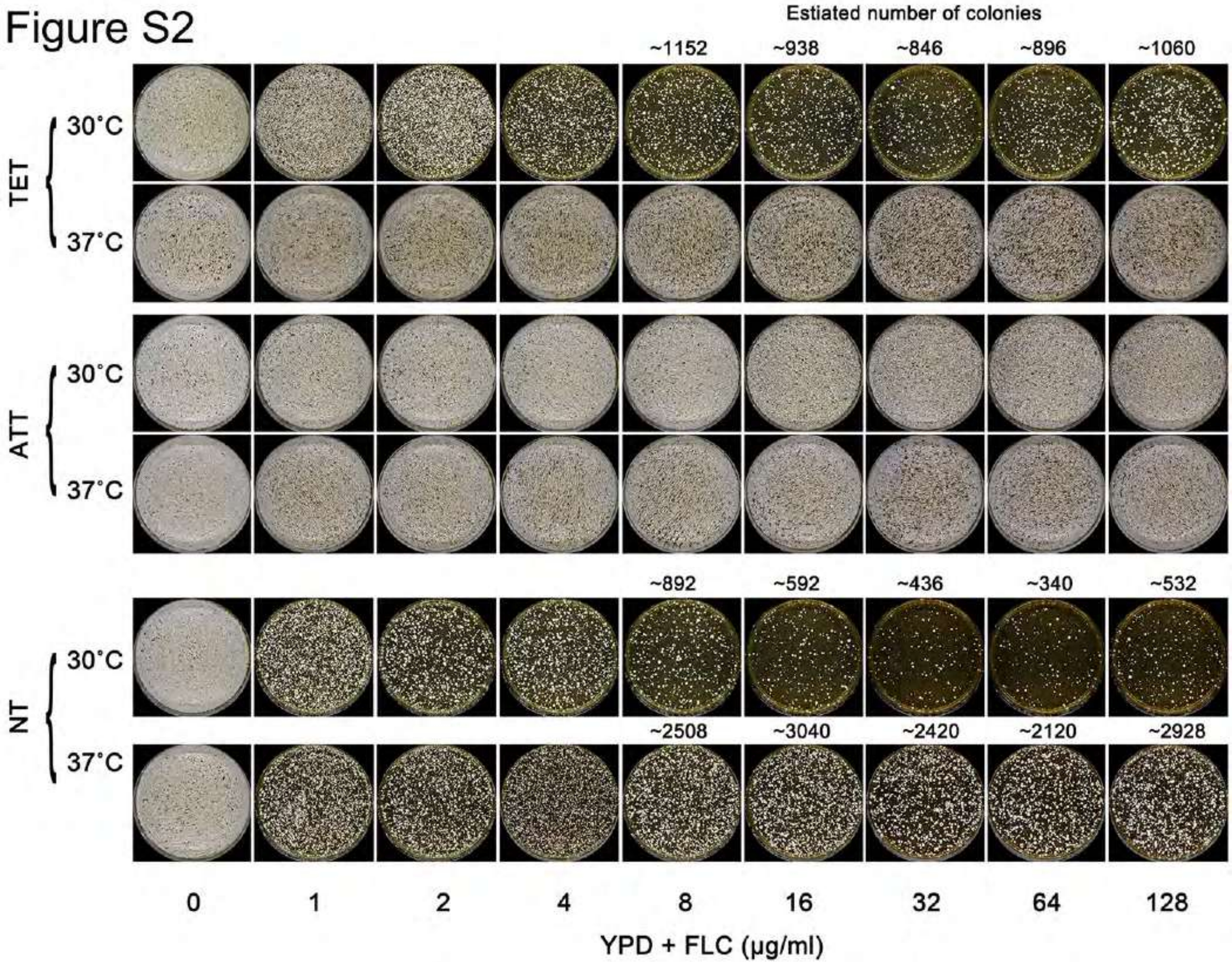
Estiated number of colonies
~1152
~938
~846
~896
~1060
TET
30°C
37°C
ATT
30°C
37°C
NT
30°C
37°C
~892
~592
~436
~340
~532
~2508
~3040
~2420
~2120
~2928
0
1
2
4
8
16
32
64
128
YPD + FLC (μg/ml)

**Figure S3**

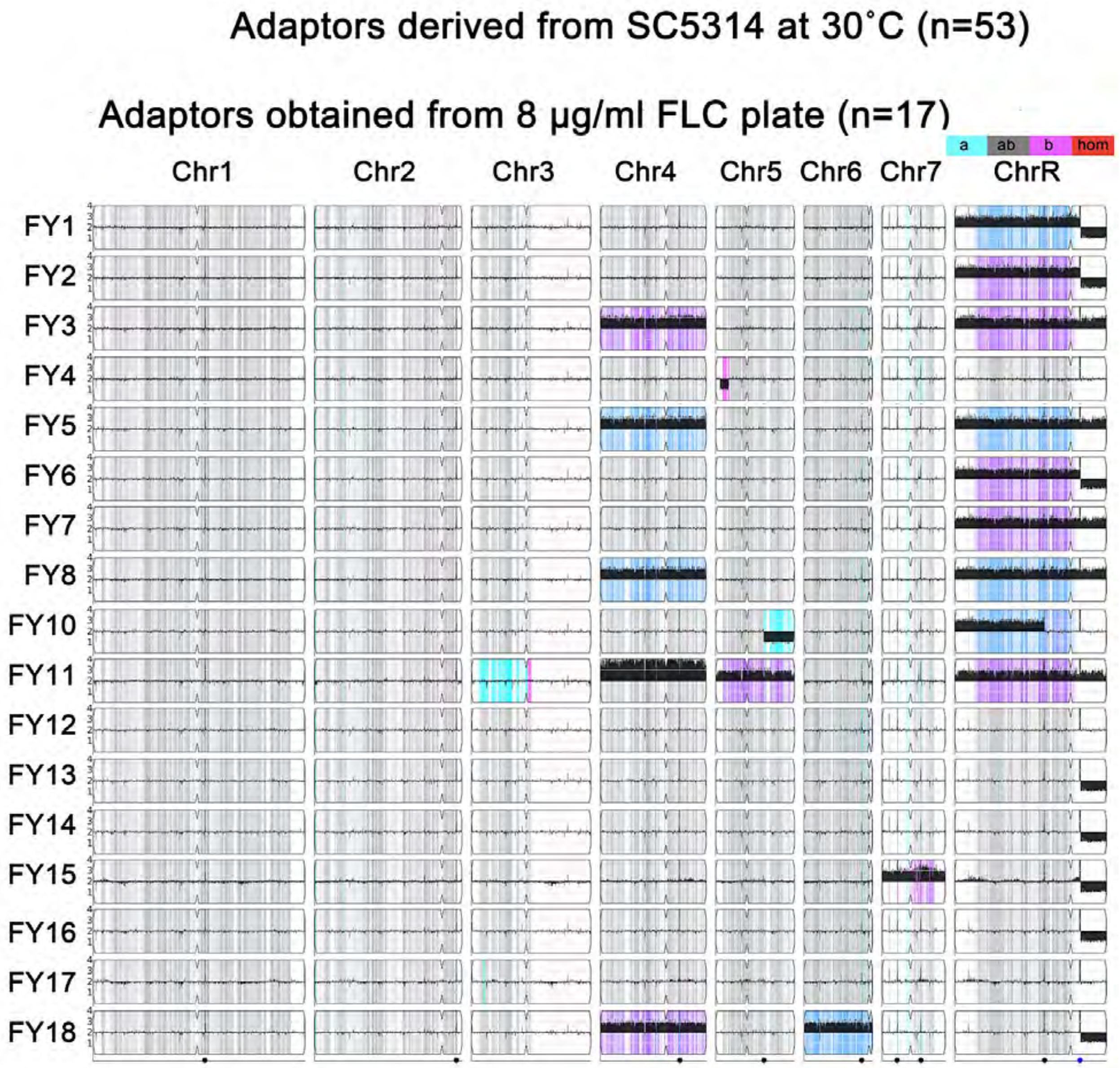

Adaptors derived from SC5314 at 30˚C (n=53)
Adaptors obtained from 8 µg/ml FLC plate (n=17)
a
ab
b
hom
Chr1
Chr2
Chr3
Chr4
Chr5
Chr6
Chr7
ChrR
FY1
FY2
FY3
FY4
FY5
FY6
FY7
FY8
FY10
FY11
FY12
FY13
FY14
FY15
FY16
FY17
FY18

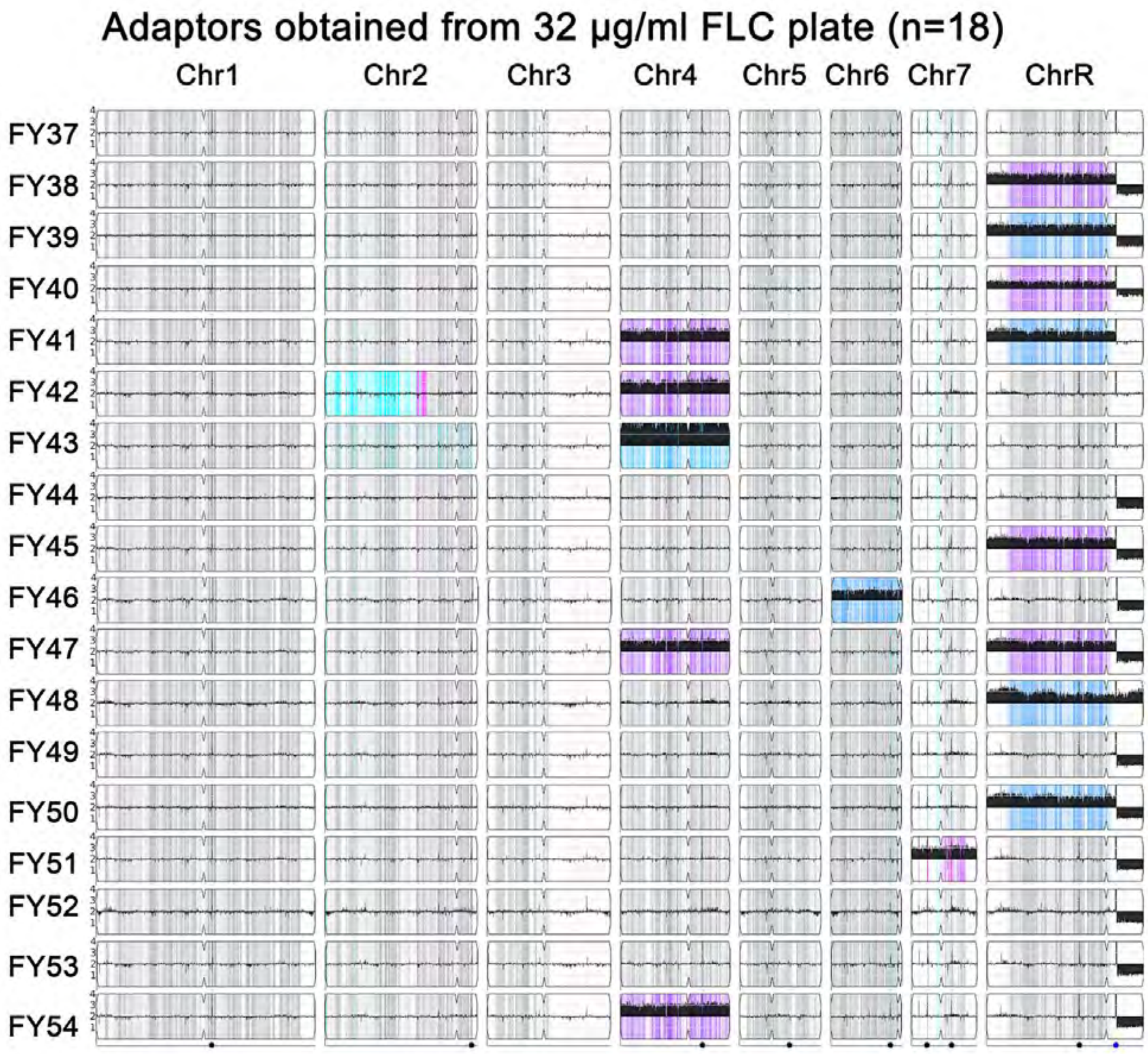

Adaptors obtained from 32 μg/ml FLC plate (n=18)
Chr1
Chr2
Chr3
Chr4
Chr5
Chr6
Chr7
ChrR
FY37
FY38
FY39
FY40
FY41
FY42
FY43
FY44
FY45
FY46
FY47
FY48
FY49
FY50
FY51
FY52
FY53
FY54

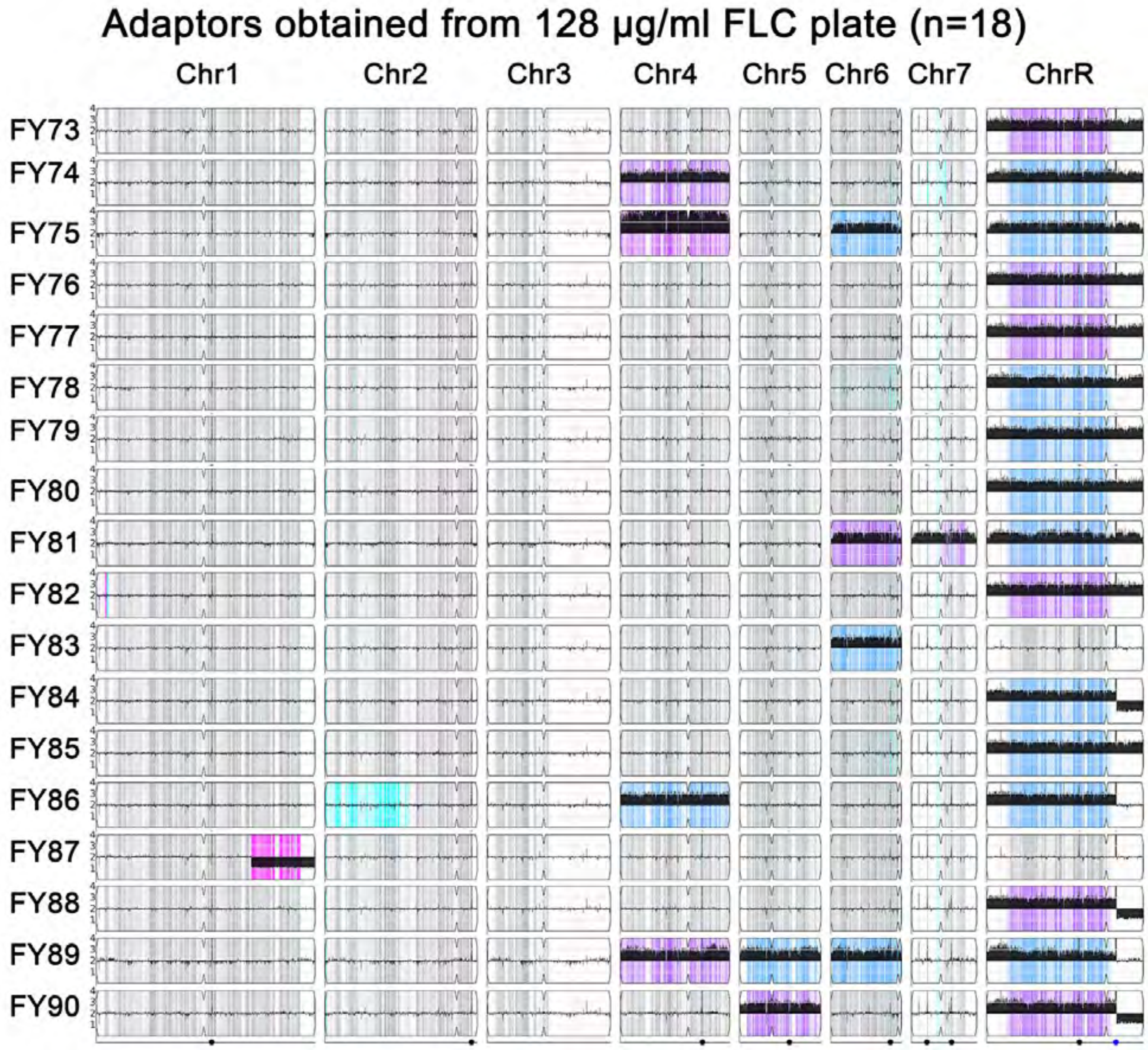

Adaptors obtained from 128 µg/ml FLC plate (n=18)
Chr1
Chr2
Chr3
Chr4
Chr5
Chr6
Chr7
ChrR
FY73
FY74
FY75
FY76
FY77
FY78
FY79
FY80
FY81
FY82
FY83
FY84
FY85
FY86
FY87
FY88
FY89
FY90

**Figure S4**

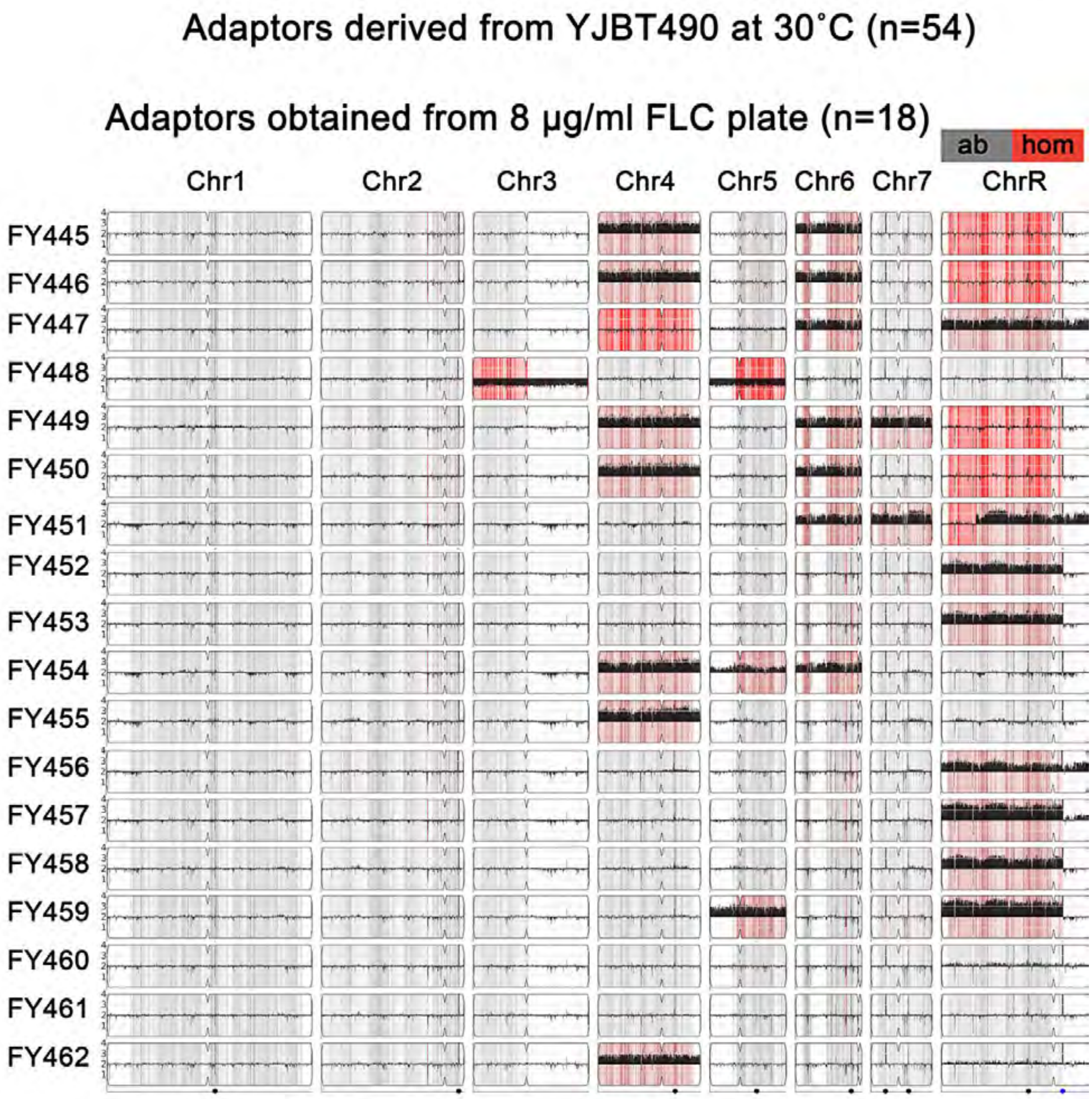
Adaptors derived from YJBT490 at 30˚C (n=54)
Adaptors obtained from 8 µg/ml FLC plate (n=18)
ab
hom
Chr1
Chr2
Chr3
Chr4
Chr5
Chr6
Chr7
ChrR
FY445
FY446
FY447
FY448
FY449
FY450
FY451
FY452
FY453
FY454
FY455
FY456
FY457
FY458
FY459
FY460
FY461
FY462

### Adaptors obtained from 32 µg/ml FLC plate (n=18)

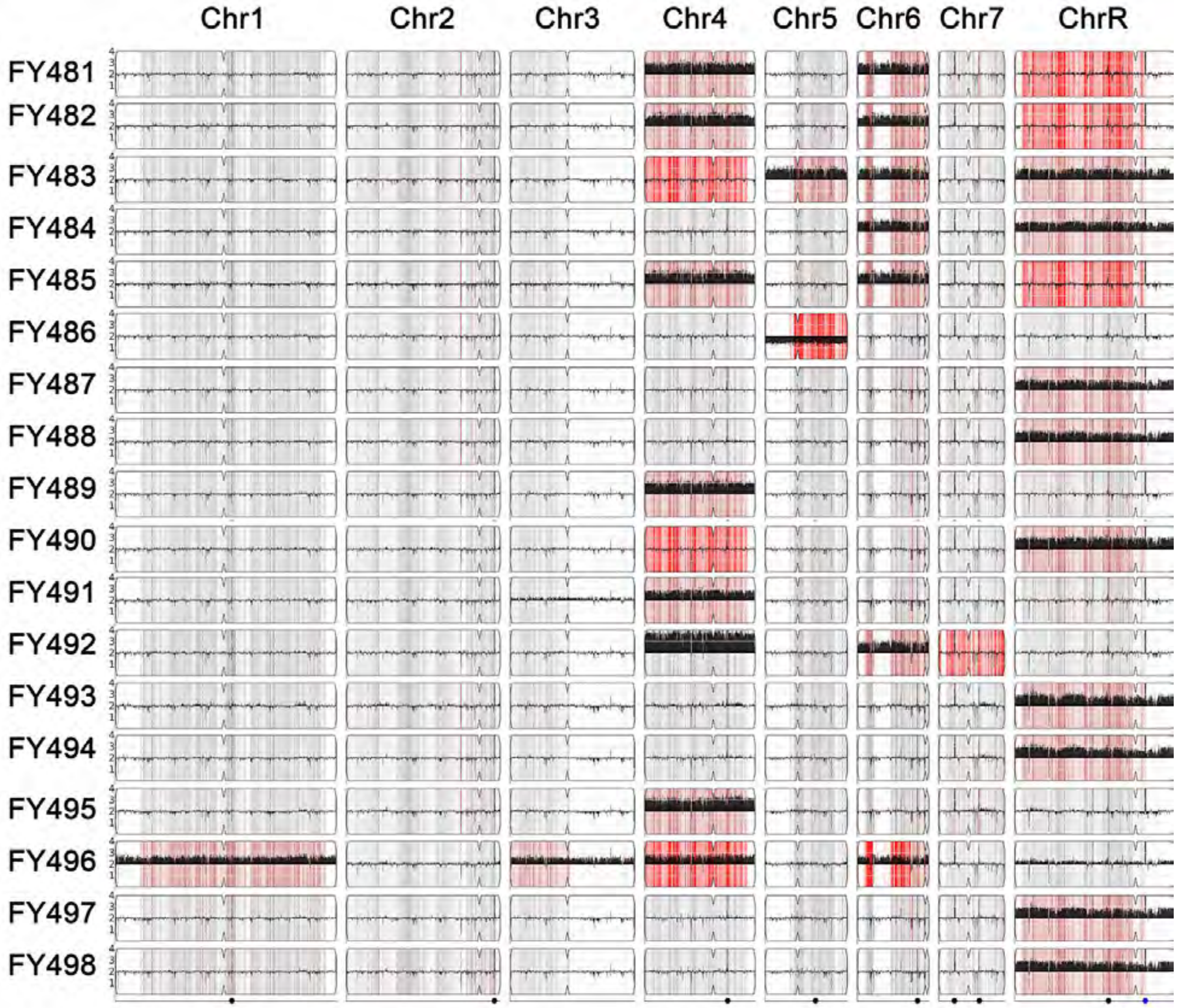

#### Adaptors obtained from 128 µg/ml FLC plate (n=18)

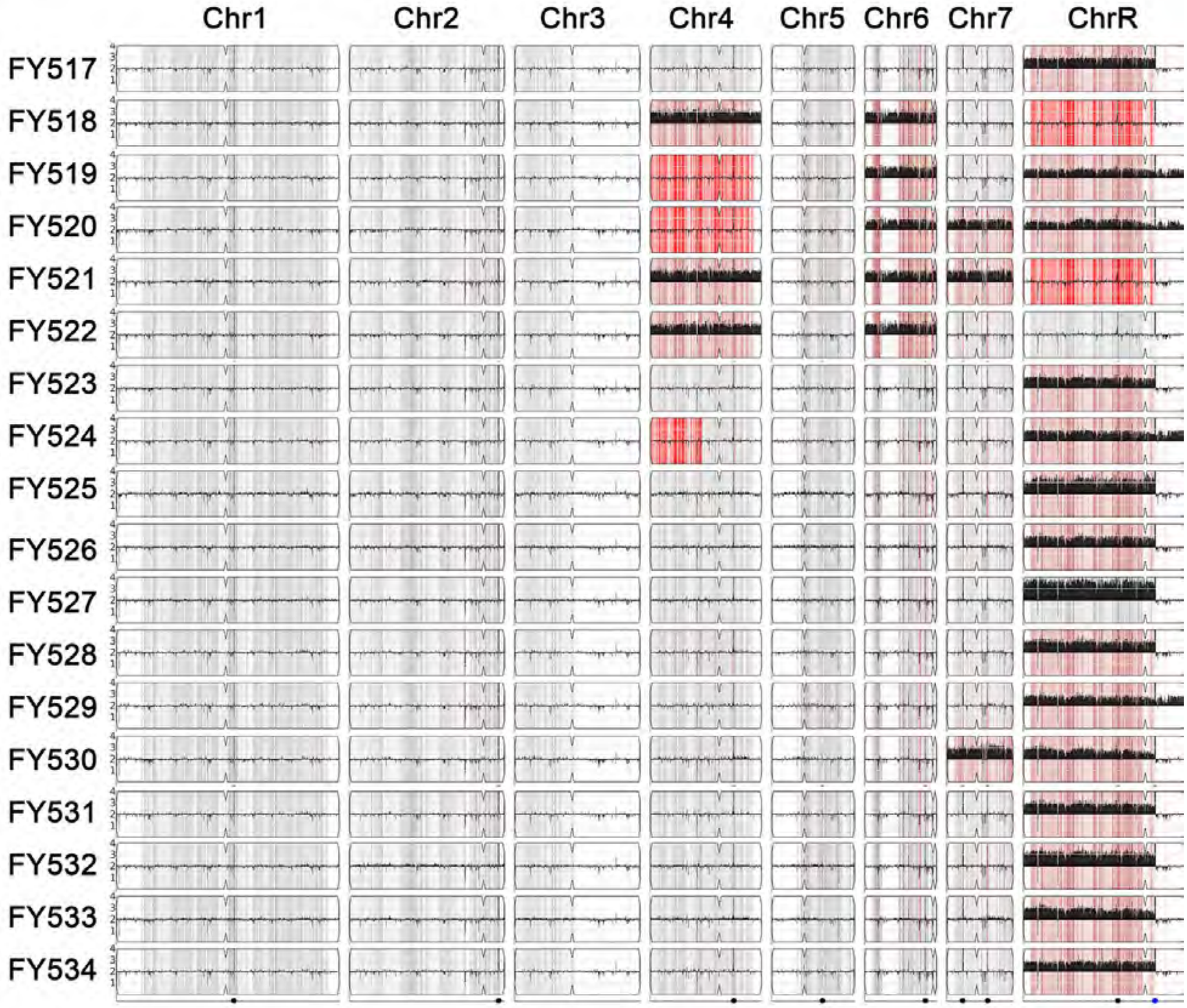

Figure S5

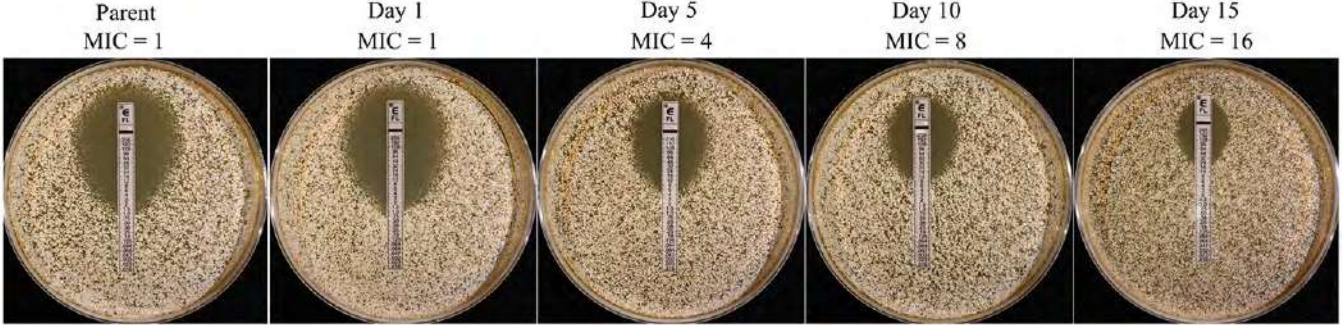

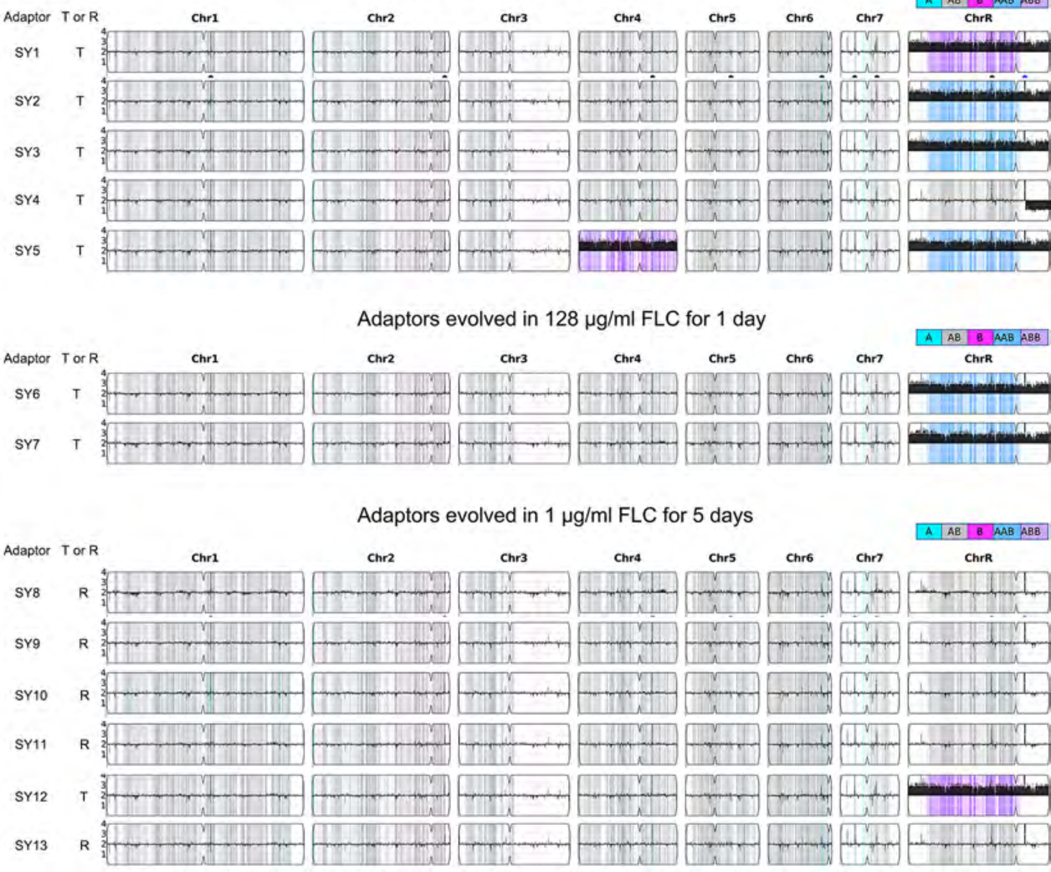
Adaptors evolved in 2 µg/ml FLC for 1 day
A AB B AAB ABB
Adaptor T or R
Chr1 Chr2 Chr3 Chr4 Chr5 Chr6 Chr7 ChrR
SY1 T
SY2 T
SY3 T
SY4 T
SY5 T
Adaptors evolved in 128 µg/ml FLC for 1 day
A AB B AAB ABB
Adaptor T or R
Chr1 Chr2 Chr3 Chr4 Chr5 Chr6 Chr7 ChrR
SY6 T
SY7 T
Adaptors evolved in 1 µg/ml FLC for 5 days
A AB B AAB ABB
Adaptor T or R
Chr1 Chr2 Chr3 Chr4 Chr5 Chr6 Chr7 ChrR
SY8 R
SY9 R
SY10 R
SY11 R
SY12 T
SY13 R

Figure S6

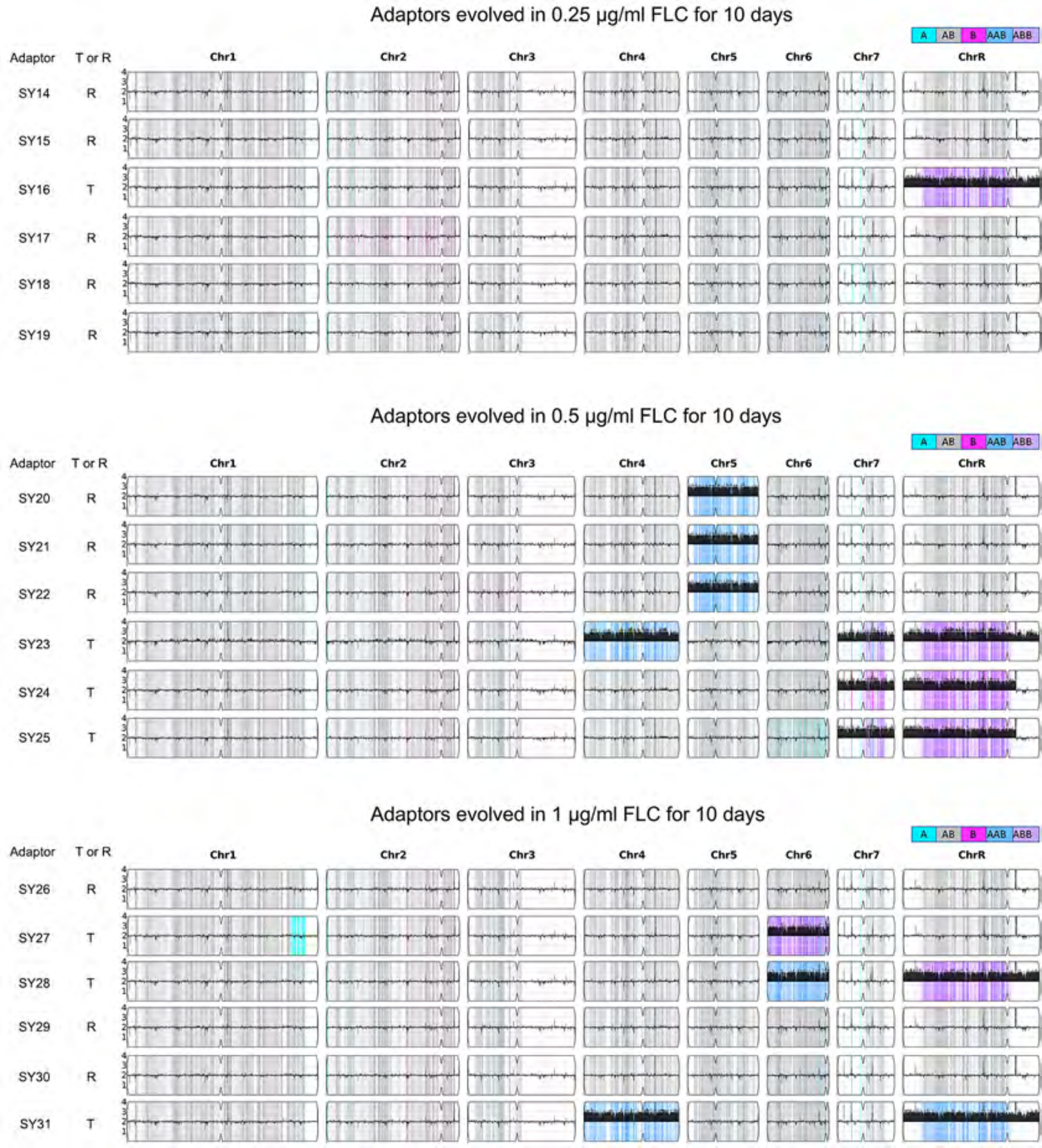
Adaptors evolved in 0.25 µg/ml FLC for 10 days
A AB B AAB ABB
Adaptor T or R
Chr1 Chr2 Chr3 Chr4 Chr5 Chr6 Chr7 ChrR
SY14 R
SY15 R
SY16 T
SY17 R
SY18 R
SY19 R
Adaptors evolved in 0.5 µg/ml FLC for 10 days
A AB B AAB ABB
Adaptor T or R
Chr1 Chr2 Chr3 Chr4 Chr5 Chr6 Chr7 ChrR
SY20 R
SY21 R
SY22 R
SY23 T
SY24 T
SY25 T
Adaptors evolved in 1 µg/ml FLC for 10 days
A AB B AAB ABB
Adaptor T or R
Chr1 Chr2 Chr3 Chr4 Chr5 Chr6 Chr7 ChrR
SY26 R
SY27 T
SY28 T
SY29 R
SY30 R
SY31 T

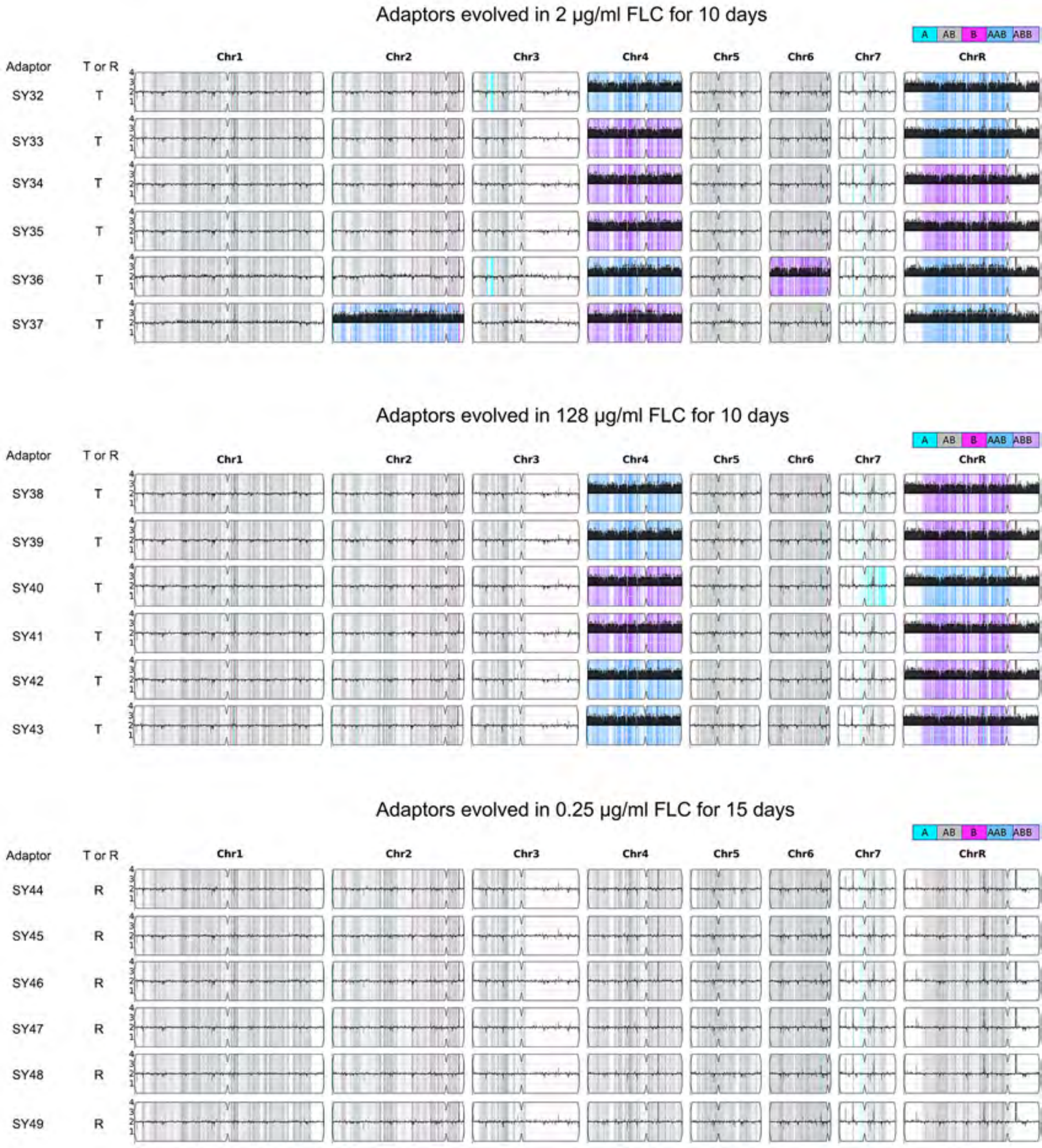
Adaptors evolved in 2 µg/ml FLC for 10 days
A AB B AAB ABB
Adaptor
T or R
Chr1 Chr2 Chr3 Chr4 Chr5 Chr6 Chr7 ChrR
SY32 T
SY33 T
SY34 T
SY35 T
SY36 T
SY37 T
Adaptors evolved in 128 µg/ml FLC for 10 days
A AB B AAB ABB
Adaptor
T or R
Chr1 Chr2 Chr3 Chr4 Chr5 Chr6 Chr7 ChrR
SY38 T
SY39 T
SY40 T
SY41 T
SY42 T
SY43 T
Adaptors evolved in 0.25 µg/ml FLC for 15 days
A AB B AAB ABB
Adaptor
T or R
Chr1 Chr2 Chr3 Chr4 Chr5 Chr6 Chr7 ChrR
SY44 R
SY45 R
SY46 R
SY47 R
SY48 R
SY49 R

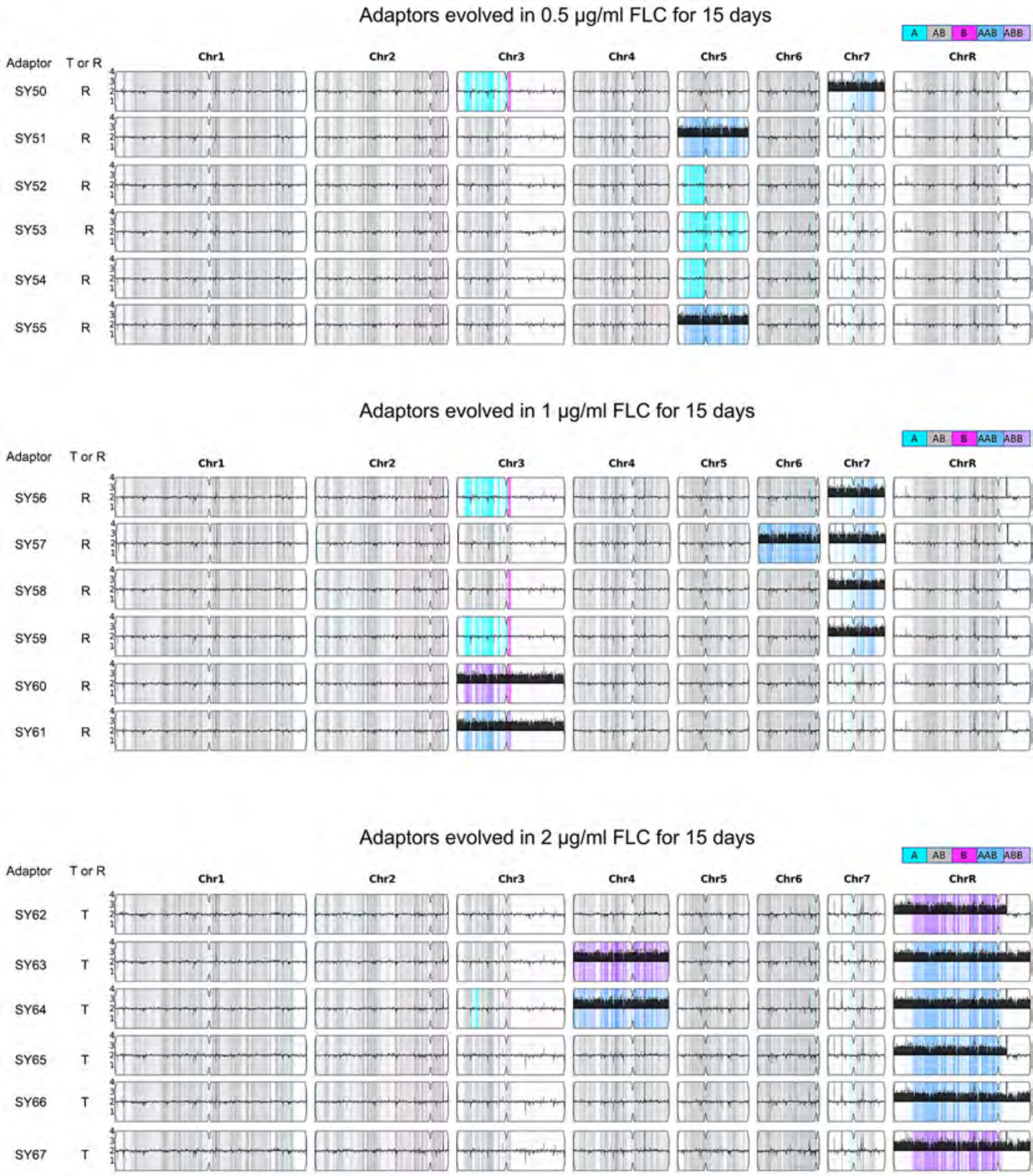
Adaptors evolved in 0.5 μg/ml FLC for 15 days
A
AB
B
AAB
ABB
Adaptor
T or R
Chr1
Chr2
Chr3
Chr4
Chr5
Chr6
Chr7
ChrR
SY50 R
SY51 R
SY52 R
SY53 R
SY54 R
SY55 R
Adaptors evolved in 1 μg/ml FLC for 15 days
A
AB
B
AAB
ABB
Adaptor
T or R
Chr1
Chr2
Chr3
Chr4
Chr5
Chr6
Chr7
ChrR
SY56 R
SY57 R
SY58 R
SY59 R
SY60 R
SY61 R
Adaptors evolved in 2 μg/ml FLC for 15 days
A
AB
B
AAB
ABB
Adaptor
T or R
Chr1
Chr2
Chr3
Chr4
Chr5
Chr6
Chr7
ChrR
SY62 T
SY63 T
SY64 T
SY65 T
SY66 T
SY67 T

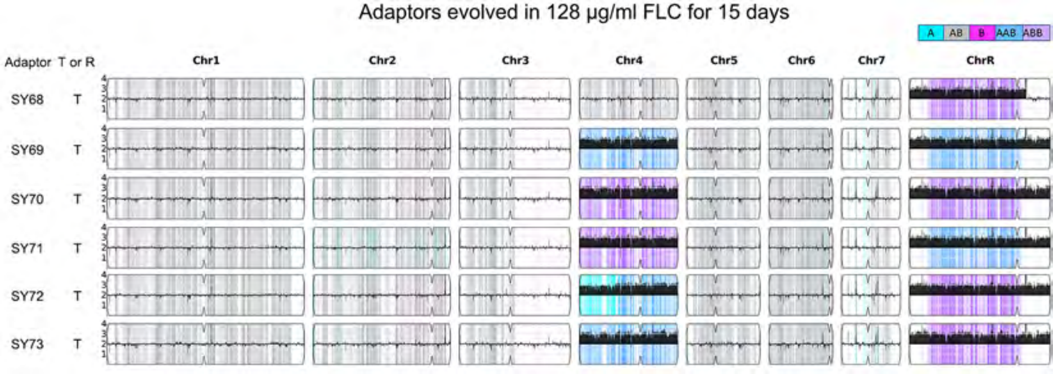

Adaptors evolved in 128 μg/ml FLC for 15 days
A
AB
B
AAB
ABB
Adaptor
T or R
Chr1
Chr2
Chr3
Chr4
Chr5
Chr6
Chr7
ChrR
SY68
SY69
SY70
SY71
SY72
SY73
T
